# Dispersal of a dominant competitor can drive multispecies coexistence in biofilms

**DOI:** 10.1101/2023.11.28.569042

**Authors:** Jacob D. Holt, Daniel Schultz, Carey D. Nadell

**Author notes:** Author for correspondence: Carey D. Nadell (ORCID 0000-0003-1751-4895)) Class of 1978 Life Science Center 78 College St. Hanover, NH 03755 USA.

## Abstract

Despite competition for both space and nutrients, bacterial species often coexist within structured, surface-attached communities termed biofilms. While these communities play important, widespread roles in ecosystems and are agents of human infection, understanding how multiple bacterial species assemble to form these communities and what physical processes underpin the composition of multispecies biofilms remains an active area of research. Using a model three-species community composed of *P. aeruginosa*, *E. coli*, and *E. faecalis*, we show with cellular scale resolution that biased dispersal of the dominant community member, *P. aeruginosa*, prevents competitive exclusion from occurring, leading to coexistence of the three species. A *P. aeruginosa bqsS* deletion mutant no longer undergoes periodic mass dispersal, leading to local competitive exclusion of *E. coli*. Introducing periodic, asymmetric dispersal behavior into minimal models parameterized by only maximal growth rate and local density supports the intuition that biased dispersal of an otherwise dominant competitor can permit coexistence generally. Colonization experiments show that WT *P. aeruginosa* is superior at colonizing new areas in comparison to *ΔbqsS P. aeruginosa*, but at the cost of decreased local competitive ability against *E. coli* and *E. faecalis*. Overall, our experiments document how one species’ modulation of a competition-dispersal-colonization trade-off can go on to influence the stability of multispecies coexistence in spatially structured ecosystems.

## Introduction

Many microbes encounter and interact with solid-liquid and air-liquid interfaces^1^, to which they may attach and embed themselves in a secreted polymer matrix to form biofilms^2–8^. This mode of growth entails fundamentally distinct elements of spatial competition and interactive behavior relative to planktonic growth^9^. Biofilm formation inherently involves a struggle for limited space and nutrients^10^, and as examples of microscopic ecosystems containing multiple strains and species, natural biofilms are expected to emerge from a mixture of mutually helpful, competitive, and actively antagonistic cell-cell sinteractions^11^. Given that space and limiting growth substrates can quickly become scarce among tightly packed multispecies groups of bacteria co-habiting a given location, how do multiple genotypes coexist despite the prevailing impact of competition^10,12–18?^ More specifically, how does the behavior of microbes in biofilm contexts influence coexistence among different microbial strains and species in manners distinct from those that occur in liquid conditions? These are core questions for the general ecology of microbial biofilm formation.

Ecological theory predicts that for *N* genotypes (different strains and species) to coexist in an ecosystem, they must be constrained by at least *N* independent limiting factors. These limiting factors can be resource types, interactions within and among species, and myriad environmental features^19–25^. When an ecosystem varies in space or time, the number of limiting factors can increase, and additional genotypes can coexist in the heterogeneous ecosystem, when otherwise they would be lost in a corresponding homogeneously mixed system^24,26^. While this theory does not necessarily predict specific mechanisms that drive coexistence for every instance, it provides an overarching framework that can aid in looking for specific mechanisms. The physiological details of biofilm formation vary substantially between different species, but most examples are heterogenous in time and space in terms of solute and matrix composition and cellular physiology^2,27,28^. Biofilm structure systematically changes through time as cells alter their metabolic activity, secrete adhesins, and many other phenotypes as groups of cells grow larger. Additionally, mechanical interactions within and among biofilm cell groups and the depletion of local resource solutes generate gradients of key resources that often feedback to population and community dynamics, and vice versa^16,29–32^.

Though we have extensive documentation of heterogeneity of biofilm architecture and cell physiology in time and space, there is at present minimal exploration of how biofilm-specific behavior influences multispecies coexistence, and how these details may be distinct from principles that operate in well-mixed liquid environments. Of the work that has studied the stability of multi-strain and multi-species coexistence in biofilms, little has measured biofilm structure at the resolution of individual cells. This cellular scale quantitative perspective can add new insight into how individual cell behavior and physiology translate to higher order group structure and composition^33,34^.

Here we developed a model biofilm community composed of *Pseudomonas aeruginosa*, *Escherichia coli*, and *Enterococcus faecalis*. All three bacterial species can behave as opportunistic pathogens and are frequently isolated from catheter associated urinary tract infections^35–37^. Our core goal was to determine how these three species assemble to form a community and what mechanisms unique to the biofilm mode of life influence the community’s steady states. To accomplish this, we use a combination of liquid shaking culture, microfluidic culture, confocal microscopy, cellular resolution image analysis, and minimal mathematical models. In well-mixed liquid culture experiments, we found that coexistence does not occur among all three species; *P. aeruginosa* displaces the other two. In biofilm culture on submerged surfaces, however, we do observe coexistence of all three species for as long as we ran the experiment (up to 25 days), with *P. aeruginosa* in the majority. We observed that *P. aeruginosa* undergoes cycles of mass dispersal that drive accompanying fluctuations in biofilm community size and cell packing architecture. By manipulating the ability of *P. aeruginosa* to disperse using a targeted gene deletion, we found that these mass dispersal events are a key driver of localized 3-species coexistence in our model biofilm system. Simple models illustrate how the observed mass-dispersal dynamics can lead to stable coexistence in a spatial setting, so long as *P. aeruginosa* is dispersing in greater proportion to its starting population size than the other two species. Dispersal-deficient *P. aeruginosa* has increased local competitive ability within multispecies biofilms and drives out *E. coli* entirely, but it is substantially deficient in its per-cell dispersal and colonization of other locations. Taken together, our results illustrate how one species’ competition/dispersal/colonization trade-off can cause temporal fluctuations in microscopic biofilm ecosystems and drive localized coexistence among different species.

## Materials and Methods

### Bacterial strains

The *P. aeruginosa* strains used in this study were derived from PA14^38,39^. The *E. coli* strains were derived from AR3110, which is a modified K-12 W3110 strain with its natural extracellular matrix production restored^40^. The fluorescent *E. faecalis* strain OG1RF and was a gift of the Dunny laboratory^41^. The fluorescent protein expression construct insertions and the deletion mutants were made here and previously using standard allelic exchange. Cultures were grown overnight in lysogeny broth (LB). Biofilm and planktonic cultures were grown in 1% tryptone broth.

### Microfluidic flow device assembly

Microfluidic chambers for biofilm culture were made with polydimethylsiloxane (PDMS) using standard soft lithography techniques. PDMS was mixed and then cured on molds of chamber sets produced using photolithography, hole-punched for inlets and outlets, and bound to #1.5 36 mm by 60 mm glass coverslips via plasma cleaning. Constant flow was generated via Harvard Apparatus Pico Plus syringe pumps loaded with either 1 mL Brandzig plastic syringes or 5 mL BD syringes. Syringes had 25-gauge needles affixed to them that were fitted with #30 Cole Parmer PTFE tubing with an inner diameter of 0.3 mm. Inlet tubing for each chamber was affixed to the media syringe, and outlet tubing was run into a petri plate for waste collection or dispersal measurements (see below).

### Competition in liquid shaking culture

Strains were grown overnight in LB medium at 37 degrees Celsius, with *P. aeruginosa* and *E. coli* being shaken at 250 rpm and *E. faecalis* being unshaken. All strains were then diluted to an OD_600_ of 0.2 in 1 % tryptone before being mixed at a 1:1 ratio. 50 µL was then transferred into 5 mL of 1% tryptone broth in a 15 mm culture tube with 2 glass beads to breakup aggregate formation. These cultures were shaken at 300 rpm. Every 24 h 50 µL was transferred into a fresh 5 mL of media. Relative abundance was measured every 24 h, immediately prior to dilution, by spotting 5 µL onto a glass coverslip and imaging 3 separate 212×212 μm fields of view.

### Biofilm culture conditions

Strains were grown overnight in LB medium at 37 degrees Celsius, with *P. aeruginosa* and *E. coli* being shaken at 250 rpm and *E. faecalis* being unshaken. Overnight cultures were diluted to an OD_600_ of 0.2 in 1% tryptone before being mixed at a 1:1 ratio. The inoculum was then transferred into planar microfluidic devices measuring 5000 µm in length, 500 µm in width and 70 µm in height. After 1 h of incubation, 1% tryptone broth was introduced to the chamber at a rate of 0.05 µl/min (corresponding to an average flow velocity of 23.4 µm/s) at 22 degrees Celsius. To measure biomass within the chamber, the biofilm was imaged by confocal microscopy at z-intervals of 0.44 µm across 3 separate 212×212 µm fields of view. BiofilmQ was then used to perform image segmentation and quantification, and Python was used to perform statistical analysis and to produce figures.

### Mean field model

To model the dynamics of a mixed population, we consider competition for resources between the different species, with dispersal triggered by high *P. aeruginosa* density. We model the growth of the different species using coupled differential equations for logistic growth while competing for resources, with *N*_*i*_ denoting the abundance of species *i*, *K*_*i*_ denoting its carrying capacity and *r*_*i*_ denoting its maximum growth rate. We have then the following system:

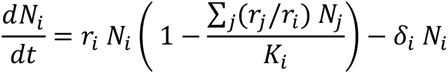

The coupling of the equations implements direct competition for resources, with the abundance of each species being limited by the abundance of the others. The coefficients for such competitive interactions are given by the relative growth between the species *r*_"_/*r*_*i*_, with the resulting dynamics favoring species that grow more quickly. Dispersal is triggered when the *P. aeruginosa* population reaches an upper threshold *N*_*up*_ and is implemented by a dispersal rate δ_*i*_. Conversely, dispersal ceases when the *P. aeruginosa* population reaches a lower threshold *N*_%&’(_, restoring δ_*i*_ = 0. We simulate the system piecewise, changing the parameter δ_*i*_ whenever the *P. aeruginosa* population crosses the upper or lower thresholds; the MATLAB ODE solver *ode113* was used to integrate the system of equations over time.

We also studied a spatial version of the model above, which implements movement of each species across a 2D surface by adding a diffusion term, arriving at a reaction-diffusion model consisting of a system of partial differential equations:

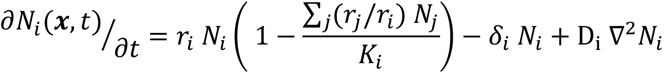

Dispersal is triggered locally when the local *P. aeruginosa* density reaches the upper threshold *N*_*up*_. We then simulate the model for an initial condition where the genotypes are randomly scattered across the surface. We use the Crank–Nicolson method, choosing time steps Δ*t* and distance increments *h* such that Δ*t* < 0.5 Δ*h*^2^/ *D* to guarantee the stability of the solution. An extended description of the model can be found in the SI Text. The MATLAB scripts used to generate all model figures are publicly available on GitHub and Zenodo. Parameters can be found in SI Table S2.

### Lattice based model

To study competition and dispersal in discrete space, we used a lattice-based model of interspecific competition. In this instance, space is represented as a 2-dimensional lattice, where each site is either vacant (state = 0) or occupied by an individual of species *i* (state *i* = 1,2,3). Simulations proceed in time steps, where each existing individual of species *i* can be removed and vacate its current site with a probability *S_i_* and give birth to another individual with a probability *β_i_*. These removal and birth probabilities are dependent on the state of the sites directly surrounding the individual. In the case of birth, a random site adjacent to the parent cell is chosen. If the chosen site is vacant, it changes to state *i*, but otherwise nothing happens and the new individual is lost^42,43^. Dispersal is triggered when the *P. aeruginosa* population within the grid reaches the *N_up_* threshold and is implemented as an increase in its removal probability *S_i_*. We note that this is an implicit implementation of cells entering the planktonic phase and subsequently being removed from the biofilm by fluid flow. The removal probability returns to baseline death rate *S_i_* once *N_up_* is reached. Interactions occur within a 250×250 grid with periodic boundary conditions. All parameters used in the simulations are listed in SI Table S2. An extended description of the model can be found in the SI Text. The Python scripts used to generate all model figures are publicly available on GitHub and Zenodo.

### Biofilm invasion assay

Cultures were grown for 4 d as described above. After 4 d of growth, the influx syringe was swapped to a new one containing one species resuspended in fresh media for 2 h. To prepare the invading strain, overnight cultures were back diluted into 50 mL of 1% tryptone and grown until mid-exponential phase. The mid-exponential phase cultures were then concentrated to an OD_600_ of 6.0. The media influent was swapped back to sterile 1% tryptone after 1 h of flow of the invading species at 0.05 µl/min. Imaging was then done at time 2 h and time 18 h post invasion. The growth of the invading strains was calculated by subtracting the invading species biovolume at 18 h from that at 2 h. The biovolume of strains was measured by imaging z-stacks of three separate 212×212 µm locations within each replicate chamber.

### Biofilm dispersal rate and colonization assay

Cultures were grown for 12 d as previously described in the biofilm culture conditions. After 12 d of growth, each biofilm was imaged, and the effluent tube was cut to 2 cm and attached to a fresh microfluidic chip to allow for colonization of a new surface. To measure the dispersal rate, the effluent tube was put into a 1.5 mL centrifuge tube with parafilm wax affixed over the top to allow for collection, while avoiding both contamination and desiccation. These samples were serially diluted, inoculated onto agar plates, and single colony forming units were distinguished by their morphology and fluorescent marker. For the colonization assay, flow into the fresh chamber ran for 2 h. For the dispersal assay, flow into the collection tube ran for 1 h to minimize growth of planktonic cells in the collected media prior to plating for CFUs. To measure the volume of colonizing cells, z-stacks of three separate 212×212 µm fields of view were acquired in the naïve chip immediately after the 2 h period of flow.

### Fluorescence microscopy

All fluorescence imaging was performed using a Zeiss 880 line-scanning confocal microscope, using a 40x/1.2 N.A. water objective. The GFP protein expressed constitutively by *E. faecalis* was excited with a 488-laser line. The mKO-κ protein that *P.* aeruginosa expresses constitutively was excited with a 543-laser line. The mKate2 protein that *E. coli* expresses constitutively was excited with a 594-laser line. All representative images were processed by constrained iterative deconvolution in ZEN blue.

### Image analysis

Native Zeiss CZI files produced by ZEN software were converted to .tiff stacks prior to being loaded into BiofilmQ, which was run using Matlab^44^. Biovolume thresholding was performed using either Otsu’s method or Robust Background with a manual sensitivity adjustment^45,46^. The segmented biovolumes for all three species were cross-checked for thresholding accuracy against the raw image data, and then dissected into a 3-D grid with cube side lengths set to 2.3 µm (giving cubes that can maximally hold ∼10 cells). The cube compartments were then used for all subsequent spatially resolved analysis. Calculated parameters were exported from BiofilmQ as .mat files and loaded into Python, where SciPy, seaborn, Pandas, NumPy, and Matplotlib were used for running statistical tests and figure generation^47–54^. For calculated cube-based parameters that require a range, we chose 10 µm as the range over which to measure (see main text for additional explanation). Operation of BiofilmQ and its array of analytical methods is described in extensive detail in Hartmann et al.^46^.

### Replication and statistics

At least four biological replicates, each defined as a single microcosm (microfluidic chamber or culture tube), were performed for each experiment across multiple weeks. For biofilm biovolume measurements, 3 regions of 212×212 µm were sampled as technical replicates within a chamber and averaged for each biological replicate. For liquid shaking culture relative abundance measurements, cells were placed between a cover slip and 3 regions of 212×212 µm were sampled and averaged for each biological replicate.

Seaborne v0.11.1 was used to generate linear regression plots. For all linear regression plots, the shaded area is the 95% confidence interval. For time resolved biofilm data, the shaded area is one standard deviation above and below the mean. Error bars denote the standard error of the mean. For box and whisker plots, the orange bar denotes the median, the box bounds denote first and third quartiles, and the whiskers denote the first and third quartiles plus 1.5 times the interquartile range. Ternary plots were either generated in python using the python-ternary library or in MATLAB using a custom script^55^. For all BiofilmQ-generated, cube-based data, mean values are weighted by biovolume for each species to account for cubes containing differing proportions of the total population. The Python scripts and corresponding datasheets used to generate figures and reported statistics are publicly available on GitHub and Zenodo.

## Results

### *P. aeruginosa*, *E. faecalis*, and *E. coli* can co-habitate over multiple weeks in biofilms

*P. aeruginosa*, *E. faecalis*, and *E. coli* were engineered to constitutively produce the fluorescent proteins mKO-κ, GFP, and mKate2, respectively, such that they could be distinguished by live-cell fluorescence microscopy. The three species were inoculated at a 1:1:1 ratio either in shaken liquid culture, or into microfluidic devices composed of PDMS chambers bonded to glass coverslips. For biofilm experiments, after a 1 h attachment period, the species were incubated under continuous flow of 1% tryptone broth medium for up to 350 h at 0.05 µL/min (average flow velocity ∼ 23.4 µm/s). As a proxy for population size, the total biovolume of each species was measured by confocal microscopy at 24 h intervals and used to calculate relative abundance. We found that the three species could cohabitate to form a biofilm community (Fig1B), which occurred despite high variance in community composition at initial timepoints (Fig1D; SI Figure S1). For liquid-culture experiments, strains were inoculated at a 1:1 ratio in identical media conditions, incubated with shaking, and transferred 1:100 to fresh medium every 24 hours. The consistent cohabitation of all three species was specific to the microfluidic biofilm environment: in our shaken liquid culture experiments *P. aeruginosa* displaced *E. coli* and *E. faecalis* from the system (Fig1A). This discrepancy in outcome between the biofilm and planktonic culture experiments suggests that the spatial constraints and/or the specific physiological states of the species in the biofilm mode of life play an important role in these three species’ ability to cohabitate within the microfluidic flow environment. The results of dual culture experiments follow the same trends, with cohabitation occurring in biofilm culture but not in planktonic culture. For the pairing of *E. coli* with *E. faecalis*, we observe cohabitation in both biofilm and planktonic culture conditions, consistent with prior work on the interaction of these two species (SI Figure S4-7)^56^.

To determine the nature of the primary ecological relationship between the three species, we measured the biovolume of each species across the first 96 h of growth under two conditions: 3-species coculture, 2-species coculture, and single species monoculture. We found that the biovolume and therefore population size of each species is higher in monoculture than in tri-culture or dual culture over this timeframe, indicating that the 3 species are primarily competing with each other in the biofilm context used here (Fig1C, SI Figure S2-S7).

Inspecting the image data in more detail reveals interesting patterns that begin to indicate what kinds of biofilm-specific behavior may be contributing to apparent coexistence in the full 3-species system. For the example time-series shown in Figure 1E, we note that the biofilm community is first dominated by *E. faecalis* and *E. coli* at early time points, followed by rapid growth of *P. aeruginosa*, which partially displaces the other two species. After reaching high densities, there is a widespread and rapid decline in *P. aeruginosa* volume and what appears to be an increase in relative abundance of the other two species. The same cycle of *P. aeruginosa* population expansion and then contraction occurs once again before the time series is halted. This pattern of *P. aeruginosa* population growth and decline was repeatable across separate runs of the experiment in a consistent manner, albeit with variation among replicates in starting conditions and the particular days on which decay and re-growth events occurred. *P. aeruginosa* dispersal behavior has been studied in considerable detail and is known to depend on rhamnolipid secretion, responses to the local environment, and responses to self^57–63^. It has previously been reported that *P. aeruginosa* monocultures grown in microfluidic chips experience rhamnolipid-dependent self-induced dispersal events, and we have recapitulated this result in our conditions (SI Figure S8)^64–66^. We speculated on the basis of these results and prior literature that *P. aeruginosa* was actively dispersing from the chambers after reaching large enough group sizes; it also appeared that the dispersal process was biased in the sense that *P. aeruginosa* departs the biofilm chambers in proportionally greater numbers relative to its resident biofilm population size in comparison with the other two species (Figure 1E). In the following section we use simple mathematical models to assess how different dispersal regimes might contribute to long-term cohabitation, and we compare modeling results with experiments to determine which of these dispersal regimes gives the best fit to our *in vitro* biofilm experiments.

**Figure 1:**
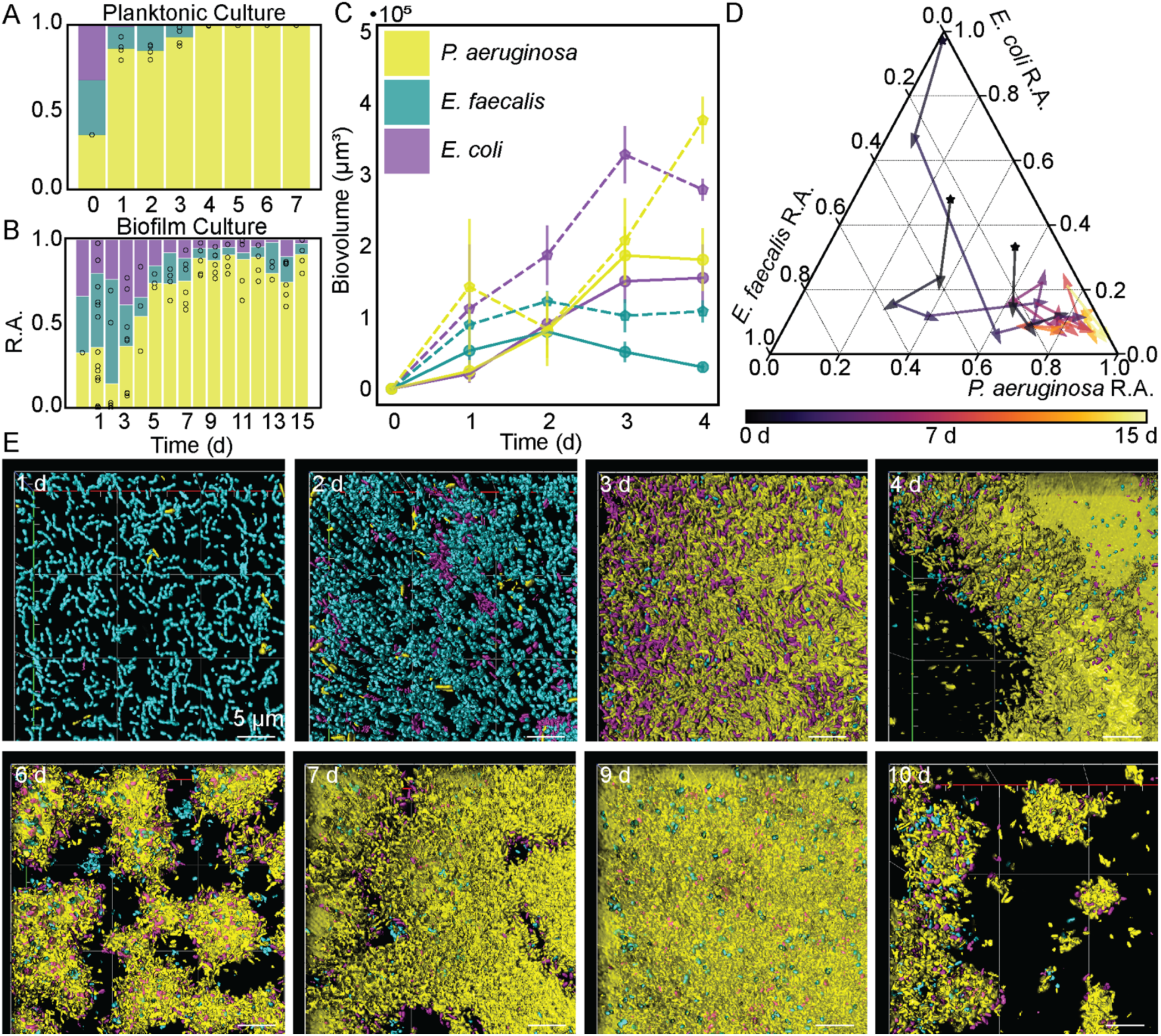
*P. aeruginosa* (yellow), *E faecalis* (cyan), and *E. coli* (purple) appear to coexist biofilm communities despite prevailing competition between them. **(A)** Relative abundances of the three species in planktonic culture (n = 4 per time point). **(B)** Relative abundances of the three species in biofilm culture (n = 3-12 per time point). **(C)** Measurement of each species’ biovolume in a 212×212 µm field of view during monoculture (dashed line) and coculture (solid line) biofilms. Dots represent the mean and error bars represent standard error (n = 3-12 per time point). **(D)** Ternary plot relative abundance trajectories from three independent biofilm replicates colored by time. **(E)** Representative 3D renderings of the three species growing in a biofilm over time (n = 3 trajectories from independent experiments).

### Density-dependent, biased dispersal of a competitively dominant species can permit co-existence with other species in the biofilm context

Our exploratory experiments above indicated that the biofilm environment can allow for cohabitation among *P. aeruginosa*, *E. coli*, and *E. faecalis*, whereas the planktonic environment does not. We also documented that dispersal events occur in biofilm environments, and that they appear to be controlled by *P. aeruginosa* and lead to preferential removal of *P. aeruginosa* from the system. We speculated that this dispersal process, by generating negative density dependence on the rate of change in *P. aeruginosa* abundance, may be particularly important as a biofilm-specific phenomenon contributing to multispecies coexistence^25,26,67–70^. Before proceeding with further analyses and experiments, we sought to verify that this idea is valid in principle using simple modeling approaches.

We explored in particular how different dispersal regimes may influence the propensity for 3 species to coexist in a local environment, parameterizing the system using the relative maximal growth rate and carrying capacity (mean local biovolume density) for each species, extracted from our experiments in the previous section (Figure 1C, SI Figure S9, SI Table S2). Specifically, we implement a growth scenario in which no dispersal occurs, another in which unbiased dispersal occurs and removes each species by equal fractions, and finally a biased dispersal condition in which *P. aeruginosa* disperses in substantially larger fraction relative to *E. coli* and *E. faecalis*.

The different dispersal scenarios noted above are based on the distinct mechanisms of biofilm dispersal regulation that have been genetically characterized in the literature^8,57,58,65,71–73^. We sought to explore unbiased dispersal because *P. aeruginosa* rhamnolipids, used as a surfactant during active dispersal, are known to be capable of dispersing biofilms formed by other species as well (Figure 2B)^64,66,74,75^. Additionally, if *E. coli* and *E. faecalis* are embedded within dispersing *P. aeruginosa* biomass, then it is conceivable that they too could be dispersed due to their spatial arrangement. Secondly, we were also interested in the case where dispersal events are biased towards removing *P. aeruginosa* (Figure 2C). This could happen if the biofilm locations that disperse are disproportionally dominated by *P. aeruginosa*, which appeared to be the case by visual inspection of our image data in the previous section (Figure 1A, Figure 2D).

**Figure 2:**
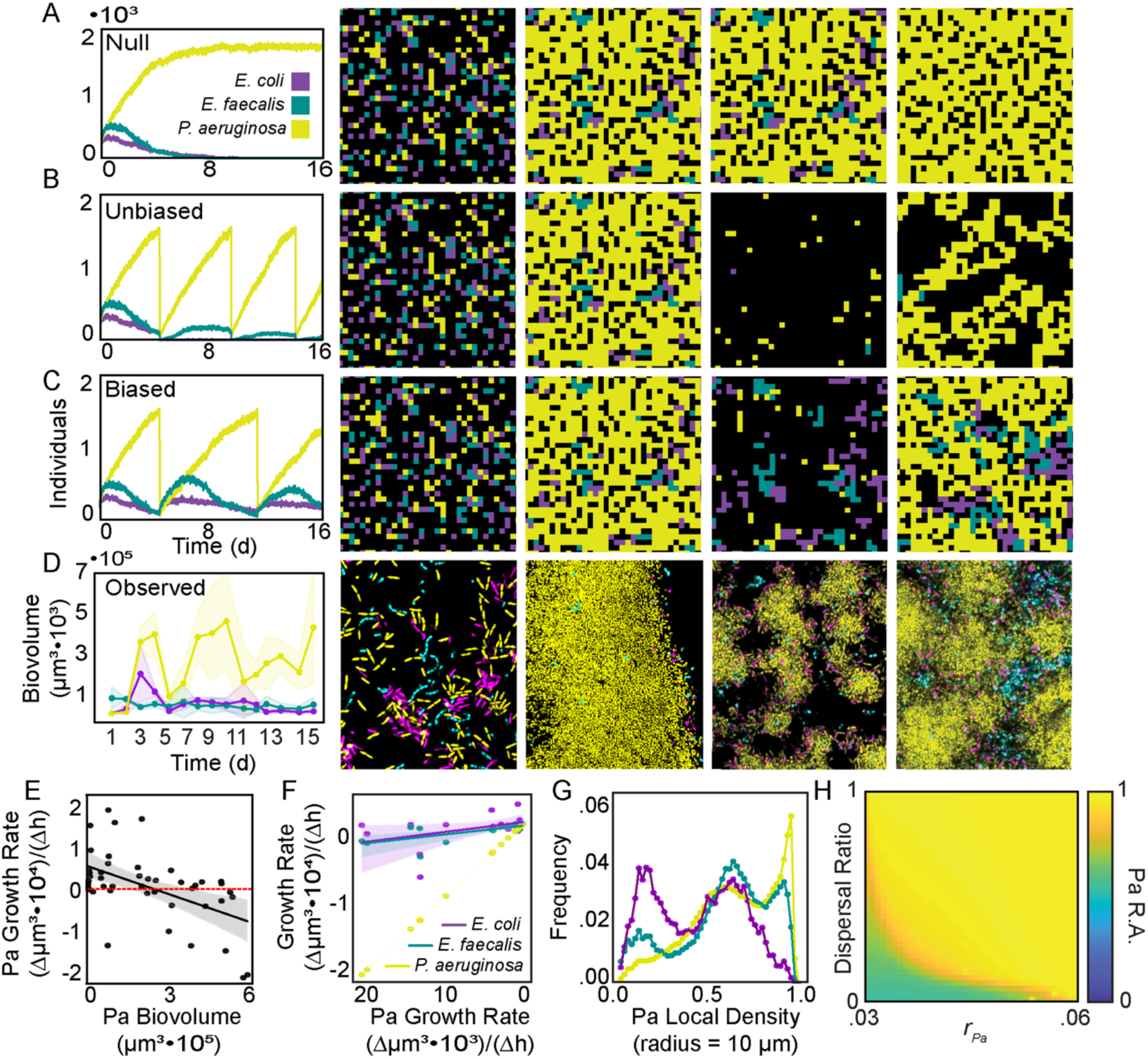
Minimal models and experimental data support dispersal as a mechanism of coexistence. **(A)** Population dynamics and images of the no-dispersal case from the stochastic-spatial model. **(B)** Population dynamics and images of the unbiased dispersal case from the stochastic-spatial model. **(C)** Population dynamics and images of the biased dispersal case from the stochastic-spatial model. **(D)** Representative population dynamics and images from the biofilm experimental dataset (One experimental run is shown here with the variance around each line derived from 3 technical replicates). **(E)** Linear regression of *P. aeruginosa* biovolume against *P. aeruginosa* growth rate (r^2^ = 0.25, p = 0.0002, n = 3). **(F)** Linear regression of negative *P. aeruginosa* growth rate against the growth rate of the other species. *E. coli* does not correlate (purple, r^2^ = 0.16, p=0.0979, n = 3). *E. faecalis* correlates moderately (cyan, r^2^ = 0.49, p = 0.0037, n = 3). **(G)** *P. aeruginosa* local density (r = 10 μm) as a frequency of total biomass (n = 4). **(H)** Phase diagram of mean-field model outcomes for equilibrium *P. aeruginosa* relative abundance as dispersal ratio (*δ_j_*/*δ_Pa_*) is varied from only *P. aeruginosa* dispersing to all species dispersing in equal proportions (dispersal ratio of 1) and *P. aeruginosa* growth rate is varied from 0.03 to 0.06.

In brief, we find that implementing biased dispersal of *P. aeruginosa* in a density-dependent manner can stabilize the community such that all three members coexist, while unbiased or no dispersal of *P. aeruginosa* cannot support all three members locally coexisting. This result is specific to the spatial stochastic model and does not hold when the stochastic-spatial system is randomly mixed at each time step (SI Figure S10D). Reproducing the system in more simplified mean-field and reaction-diffusion models gives qualitatively similar results to the stochastic-spatial model: biased dispersal of *P. aeruginosa* can stabilize the community such that all three members coexist, while unbiased or no dispersal of *P. aeruginosa* cannot support all three members locally coexisting (SI Figure S10, S11). By varying the dispersal ratio from biased (ratio of 0) to unbiased (ratio of 1) and varying the fitness of *P. aeruginosa* from low to high in the mean-field model, we can visualize the relationship between *P. aeruginosa* relative fitness and the amount of dispersal bias needed to maintain coexistence (Figure 2H).

We note that our results here are distinct from the well-explored connection between coexistence in multi-patch systems through a competition-colonization trade-off, where all species experience a strict negative relationship between local competitive ability and dispersal/colonization ability^25,76–82^. Our modeling work here focuses on and supports the simple intuition that one competitively dominant species, if it regularly disperses without carrying significant amounts of the other two species with it, can permit stable coexistence with other less locally competitive species^25,67,68,83^. This occurs because the dispersal events selectively lower the time-averaged abundance of the stronger competitor, giving the other species space and time to replicate sufficiently to maintain a positive population size.

The modeling results and experimental observations above led us to speculate that when dispersal events occur in long-lived biofilms of *P. aeruginosa*, *E. coli*, and *E. faecalis*, these events are driven by *P. aeruginosa* and remove them from the system in greater proportion to their resident biofilm population size. We directly tested this idea by quantifying the population dynamics of each species in relation to changes in *P. aeruginosa* local abundance within the 3-species system. Using the time course data from our 350 h experiments, we calculated the change in each species’ biovolume – which again here is a proxy for population size – as a function of the change in the biovolume of only *P. aeruginosa* in the time following dispersal events. This analysis was consistent with our hypothesis that *P. aeruginosa* dispersal events are primarily removing *P. aeruginosa* from the system. As anticipated, the *P. aeruginosa* time-averaged rate of population change is positive when they are at low density and negative when they are at high density, and this trend is similar both in monoculture and co-culture (Figure 2E, SI Figure S13B). This latter point emphasizes that *P. aeruginosa* is driving its own dispersal dynamics mostly or completely independently of the presence of the other two species. Additionally, the per day changes in abundance of *E. coli* and *E. faecalis* are only weakly correlated with the per day change in abundance of *P. aeruginosa*. This quantification reinforces our intuition from the previous section and suggests that during mass dispersal events, it is primarily *P. aeruginosa* that is departing the system (Figure 2F, SI Figure S13A).

We did note that the correlation between declines in *E. faecalis* and *P. aeruginosa* was statistically significant if minor in terms of magnitude. We wondered why this might be the case and began by exploring the spatial relationship between the three species. We noted that while *E. coli* and *E. faecalis* tended to be found around the periphery of *P. aeruginosa* cell groups, only *E. faecalis* could also be found in the middle of *P. aeruginosa* cell groups. We could confirm this visual intuition quantitatively by measuring the spatial occurrence of *E. coli* and *E. faecalis* in relation to the local density of *P. aeruginosa*, as well as by calculating the distance to the closest neighbor for each species-pairing (Figure 2G, SI Figure S14). These observations are consistent with the explanation that the portion of *E. faecalis* population found within *P. aeruginosa* cell groups is driving the correlation between declines in *P. aeruginosa* and *E. faecalis*.

All told, our data support the interpretation that mass dispersal events are driven primarily by *P. aeruginosa* in a density-dependent manner, and that because of the spatial relationships among the three species, *E. coli* and *E. faecalis* are less likely to depart the system during these dispersal events, relative to the fraction of the community they occupy. In the following sections we explore in greater detail the correspondence between our experimental system and the abstraction of the modeling work, and we consider the effects of dispersal behavior on competition at larger spatial scales involving multiple resource patches.

### A *P. aeruginosa* mutant with reduced dispersal does not permit local 3-species co-existence

To assess the idea that cyclic dispersal of *P. aeruginosa* is necessary for the persistent cohabitation observed in our three-species biofilm community model, we generated a *P. aeruginosa* strain with reduced dispersal relative to the parental wild type PA14. Specifically, we produced a strain harboring a clean deletion of *bqsS*, which encodes the transmembrane sensor of the BqsR/BqsS two component system that responds to Fe(II) concentration^84^. One of the primary phenotypes of a *bqsS* deletion mutant is strongly reduced biofilm dispersal, caused at least in part by a significant decrease in production of surfactants and proteases^62,85,86^. Thus this mutant afforded an opportunity to test the effects of reduced biofilm dispersal by *P. aeruginosa* on multispecies biofilm community dynamics in co-culture with *E. coli* and *E. faecalis*.

We first took note of the obvious architectural differences between biofilms containing WT *P. aeruginosa* in comparison with the Δ*bqsS* derivative. While WT *P. aeruginosa* biofilms are heterogenous and harbor regions with lower local density, Δ*bqsS* biofilms are skewed towards higher local density (SI Figure 14A). This architectural difference was maintained in the 3-species biofilms, where WT PA14 displayed regions of high density *P. aeruginosa* (which permitted little of the other two species to enter), and other, lower-density regions in which all three species were observed (Figure 3A left). Biofilms containing Δ*bqsS P. aeruginosa* appeared to contain almost exclusively high density regions of the Δ*bqsS* mutant, with few detectable cells of the other species (Figure 3A right; SI Figure 15-17). Quantification of relative abundance of *P. aeruginosa* in the two conditions confirmed these observations: the Δ*bqsS* mutant rose to nearly 100% of the community in most locations and replicates of the experiment, while WT *P. aeruginosa* stabilized at ∼85% as observed above (Figure 3B). Notably, averaged over all runs of the experiment, the Δ*bqsS* deletion mutant accumulated double the total biovolume over the course of biofilm growth relative to the wild type PA14 parental strain. These data are all consistent with the hypothesis that the Δ*bqsS* mutant disperses less than its parental strain, and that its decreased or eliminated dispersal reduces the ability of the three species to cohabit a patch of biofilm growth on the sub-millimeter scale.

**Figure 3:**
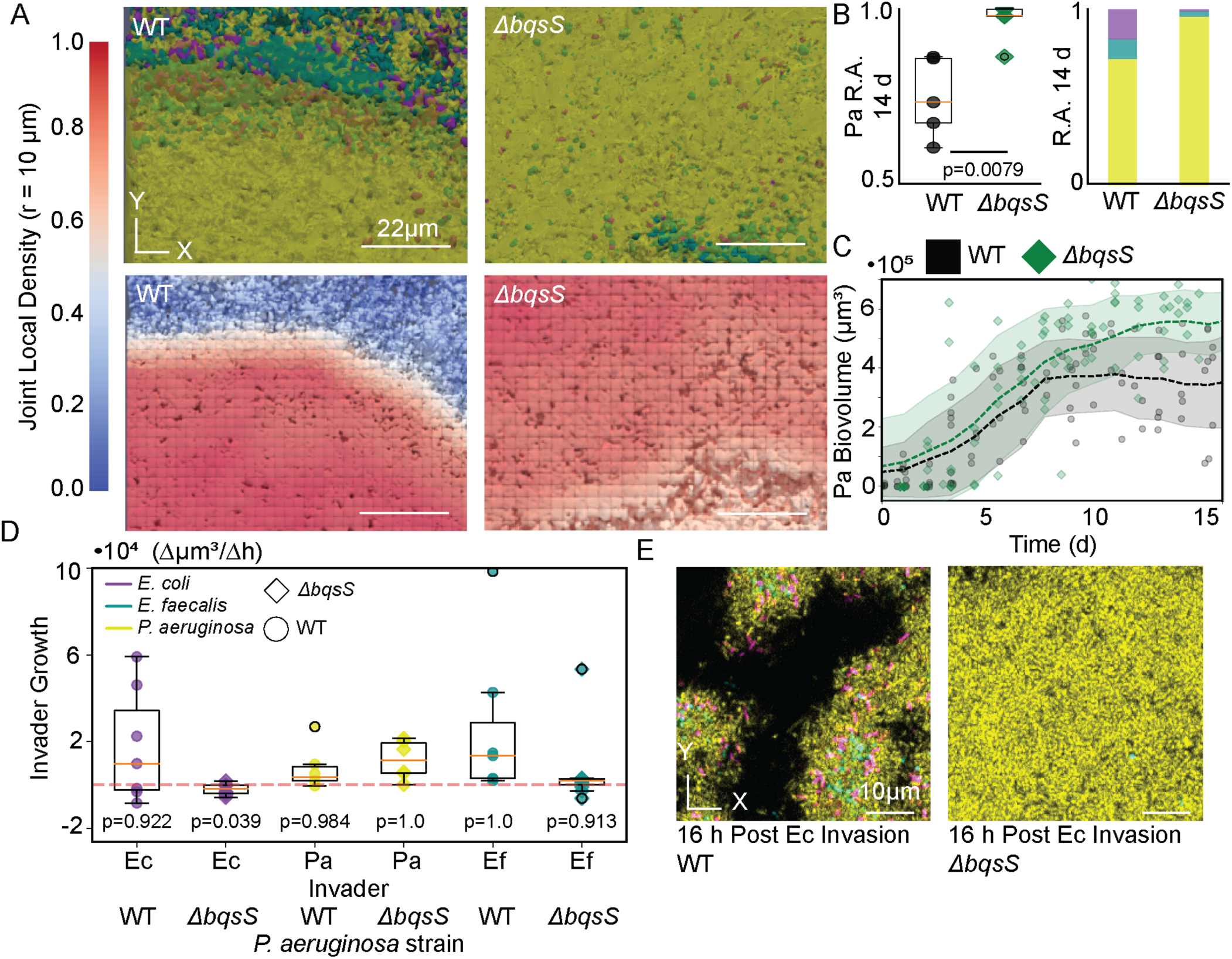
*P. aeruginosa ΔbqsS* has increased biofilm local density and invasion resistance. **(A)** 3D renderings of *P. aeruginosa* WT and Δ*bqsS* biofilms and corresponding joint local population density measurement (r = 10 μm). *P. aeruginosa* is shown in yellow, *E. coli* is shown in purple, and *E. faecalis* is shown in cyan. **(B)** Box and whisker plots showing that *P. aeruginosa* WT has significantly lower relative abundance in the three-species community relative to the *ΔbqsS* mutant (Mann-Whitney U-test, n = 5). **(C)** Time-averaged trajectories of WT *P. aeruginosa* and *ΔbqsS* biovolume in the biofilm community. Dashed lines represent the time-averaged mean (3 d window) and the shaded region represents one standard deviation above and below the time-averaged mean (n=3-10). **(D)** Box and whisker plots of single species growth after invasion into a two species biofilm showing that *E. coli* cannot invade the *ΔbqsS*-*E. faecalis* biofilm (Wilxocon tests against null that data fall above zero, n = 6-10). **(E)** Representative images of the invasion of *E. coli* into a 96 h biofilm composed of *E. faecalis* and *P. aeruginosa*.

A useful criterion for coexistence of a given set of species is that they can each increase in abundance when rare in the community^25^. An effective test of this criterion is to determine if each species can invade the community, that is, propagate to stable positive population size from initial rarity^25,87,88^. Thus far we have been measuring the population dynamics of *P. aeruginosa*, *E. coli*, and *E. faecalis* from 1:1:1 starting inoculations. To conclude this section, we sought to test the strict coexistence criterion that each species can invade biofilms composed of the other two. We did this by growing 2-species biofilms for each potential combination of the 3 species for 96 hours, followed by a pulse of the third ‘invader’ species over the course of 2 hours. This full set of invasion experiments was performed with both the wild type PA14 background and the Δ*bqsS P. aeruginosa* deletion mutant. Figure 3D illustrates that each of the 3 species, on average, registers positive and stable population sizes after colonizing a pre-existing biofilm of the other two species when *P. aeruginosa* is wild type (Figure 3E, SI Figure 18). With the Δ*bqsS* strain, *E. faecalis* retains the ability to invade, but now *E. coli* cannot, illustrating that the strongly reduced dispersal phenotype of the Δ*bqsS* strain interferes with coexistence of the three species (Figure 2D).

### The local competitive advantage of *P. aeruginosa ΔbqsS* trades off against the ability to disperse to other locations

The results reported above indicate that the repeated mass dispersal behavior of wild type *P. aeruginosa* PA14 decreases their time-averaged abundance on a local scale and opens space sufficiently often to permit localized coexistence with *E. coli* and *E. faecalis*. The Δ*bqsS* deletion mutant of *P. aeruginosa*, whose dispersal is substantially reduced, does permit coexistence with *E. faecalis*, albeit with *E. faecalis* at very low population size, but not with *E. coli*, which is no longer able to invade a resident biofilm of the two other species. We hypothesized that wild type *P. aeruginosa*, though not able to displace the other two species locally, may be balancing a trade-off between local competition and dispersal to other locations relative to the growth behavior of a Δ*bqsS* mutant.

We sought to test this possibility by comparing the relative local population growth and dispersal of wild type versus Δ*bqsS* PA14, noting that the Δ*bqsS* mutant does accumulate greater biomass locally than the wild type strain (Figure 4A). When the effluent from chambers containing these two different strains of *P. aeruginosa* (growing in co-culture with *E. coli* and *E. faecalis*) was used to colonize new microfluidic devices, however, the wild type strain of PA14 showed substantially higher colonization, including deposition of large groups of cells, which had either been released from the upstream chamber as dispersing cell groups or formed by aggregation after arriving in the new chamber. This rapid formation of three-dimensional biofilm structure is likely advantageous in environments containing exogenous threats such as bacteriophage and predators^89–95^. Additionally, these cell groups were consistent in spatial arrangement and composition of the three species within the original upstream biofilms containing WT *P. aeruginosa*, suggesting a continuation of coexistence across patches. The Δ*bqsS* mutant, by contrast, was slower to colonize downstream chambers, consistent with the idea that its local competitive advantage carried a cost of decreased ability to disperse and colonize other locations distal to the focal patch in which biofilm growth was occurring (Figure 4B-D).

**Figure 4:**
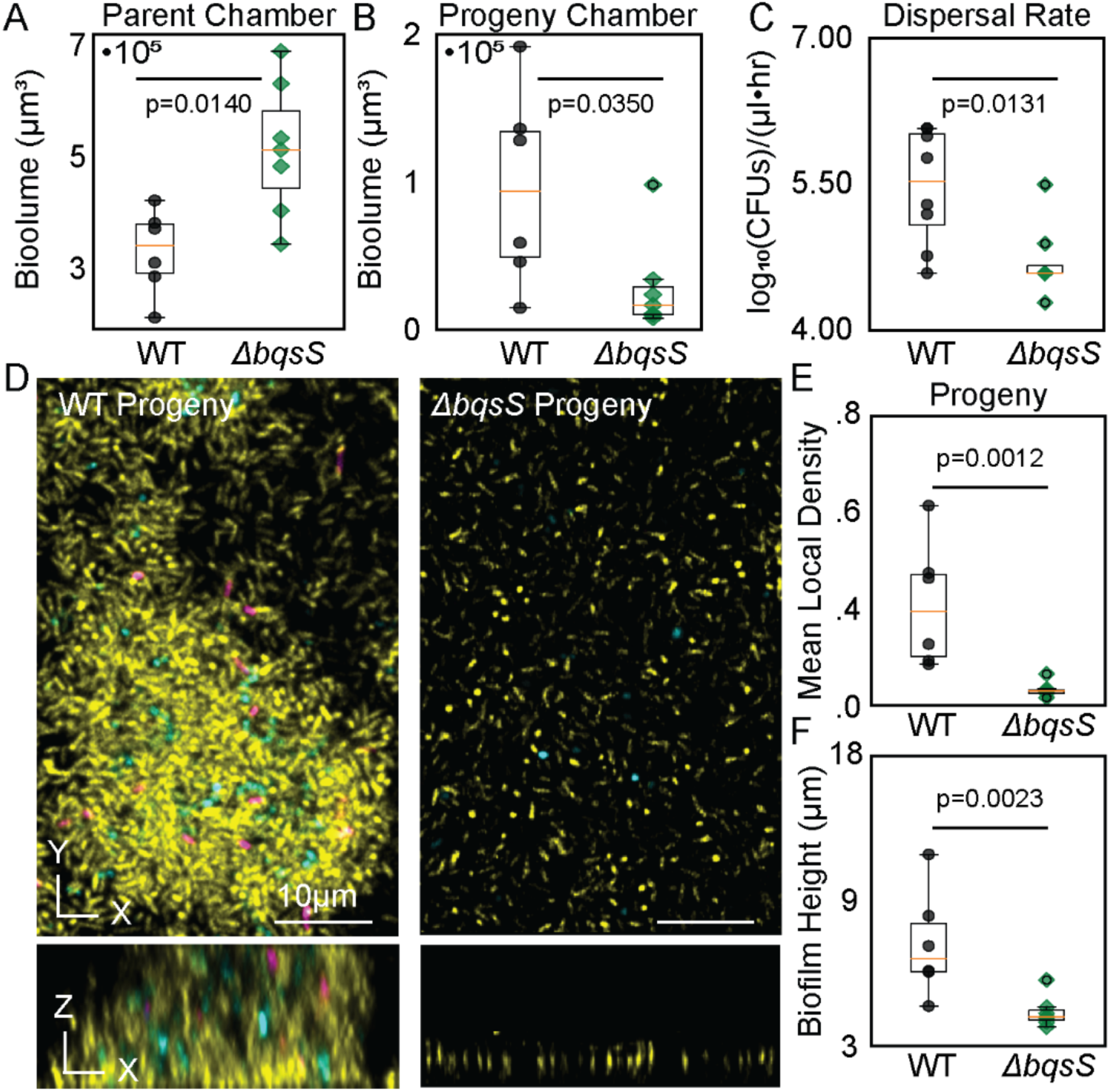
A *P. aeruginosa* BqsS deletion mutant has a lower dispersal rate and decreased colonization ability in comparison with *P. aeruginosa* WT. **(A)** Box and whisker plots showing significantly higher biovolume in the *ΔbqsS P. aeruginosa* parent/seeding chamber relative to WT (Mann-Whitney U test, n = 6). **(B)** Box and whisker plot showing significantly less *ΔbqsS* biovolume in the colonized chamber relative to WT (Mann-Whitney U test, n = 6 chambers). **(C)** Box and whisker plots showing a significant difference in CFU dispersal rates between WT and *ΔbqsS* (Mann-Whitney U test, n = 7). **(D)** Representative images of WT and *ΔbqsS* colonizing a sterile progeny chamber. *P. aeruginosa* is shown in yellow, *E. coli* is shown in purple, and *E. faecalis* is shown in cyan. **(E)** Box and whisker plots showing that the WT, mean, joint, local density (r = 10 µm) is significantly higher within the newly colonized chamber (Mann-Whitney U test, n = 6). **(F)** Box and whisker plots showing that the biofilm height (µm) of WT is significantly higher within the newly colonized chamber (Mann-Whitney U test, n = 6).

## Discussion

Using microfluidic culture methods, we generated a model three-species biofilm community composed of *E. coli*, *E. faecalis*, and *P. aeruginosa*, which we found can coexist with each other over weeks-long time scales. Coexistence of the three species, each of which can invade the other two from rarity to a positive population size, is specific to biofilm culture conditions: in shaken liquid culture, *P. aeruginosa* competitively displaces the other two species. We found that 3-species coexistence depends on cyclic mass dispersal that is primarily driven by *P. aeruginosa*, which recapitulates similar cycling of dispersal and re-growth in biofilm culture on its own. Via this pattern of repeated dispersal, *P. aeruginosa* lowers its time-averaged abundance and generates variance in proportion of space occupancy over time, which is a key factor allowing the other two species to coexist with it on a local scale. A Δ*bqsS* mutant of *P. aeruginosa*, whose dispersal is far reduced relative to the parental PA14, permits cohabitation with *E. faecalis,* but no longer with *E. coli*. However, the same mutant has considerably lower ability to colonize new locations elsewhere relative to wild type *P. aeruginosa*. These results highlight the ability of biofilm dispersal/competition trade-offs to shape microbial community composition across single and multi-patch spatial scales.

Our experimental results are consistent with long-standing ecological theory establishing that both temporal and spatial heterogeneity can create additional factors capable of limiting different species’ abundance^24,26^. Using detailed image analysis, we find that in our experimental system, *P. aeruginosa* dispersal creates a heterogenous biofilm architecture that underlies *E. coli*’s ability to coexist in the community. Numerous physical and biological mechanisms through which the biofilm mode of life generates variation among the cells dwelling within have been identified and characterized^30,96^. However, the extent to which these biofilm specific features generate additional limiting factors pertinent to multispecies coexistence, allowing biofilms to support greater numbers of species than well-mixed counterparts, has been relatively underexplored^97,98^.

Here, we have shown in detail that one specific type of biofilm behavior, active dispersal of a single, dominant species, can increase the number of species that a biofilm can maintain by decreasing the dispersing species’ time-averaged rate of change in population size and generating both spatial and temporal heterogeneity in the local biofilm ecosystem. Density-dependent dispersal is just one of many forms that dispersal from a biofilm can take, it should be noted. Dispersal from a biofilm can occur as a passive result of physical forces, an active response to environmental signals, or as an active response to self-created signals^6,8,27,28,58,71,72^. Whole or subsections of biofilm growing under shear stress can be removed en masse in what are often referred to as sloughing events. Less drastically, the growing edge of a biofilm often loses population members continuously, though in lower numbers relative to sloughing events, in an erosion-like process^99^. The environmental cues that drive dispersal are diverse and vary between different species, but quorum sensing regulation and starvation response are commonly involved^72,73,100–103^. How biofilms’ often high variance in structure over time and space impacts microbial ecosystem species richness remains an important direction for future work using both in vitro systems accessible to live time lapse microscopy, as well as more naturalistic systems and settings^104–106^.

Though we have identified how a cyclic dispersal process can contribute to multispecies coexistence, we cannot say for sure how other types of cell-cell interaction may underlie some of the fundamental competitive dynamics in the three species system studied here. We did not, for example, examine the potential roles of secreted substance-based interactions, which likely (at least in part) account for the competitive dominance of *P. aeruginosa* in the liquid culture condition and its ability to occupy the majority of the biofilm community even when it is dispersing regularly. For example, PA14 produces copious biofilm matrix including polysaccharide, protein, and DNA components, which explains their robust biofilm growth and has been shown to contribute to competition for space directly^107–110^. *P. aeruginosa* also produces a broad array of molecular weapons – such as pyocyanin, hydrogen cyanide, tailocins, and contact-dependent Type VI Secretion Systems, among others – all of which have been explored in some depth previously ^111–116^. Some of these factors may well contribute to the baseline competitive dynamics in our experiments here, though we note that the constant flow used in our biofilm culture conditions can drastically shrink the range of influence of diffusible secreted compounds^117^. On the other hand, prior work has indicated that *E. coli* and *E. faecalis* can interact synergistically under some conditions^56,118^ – *E. coli* chemotaxes to autoinducer-2 released by *E. faecalis*, for example, which can promote their co-aggregation in biofilms^56^. But we would emphasize again here that the overall primary ecological interaction we could register in all 2-species and 3-species conditions was competition, in which all community members’ population sizes were reduced relative to monoculture conditions. Biofilms, like all ecosystems, emerge from a complex interplay of mutually helpful, competitive, and actively antagonistic interactions among their constituent members; fully disentangling how these forces combine to yield community structure, richness, and stability remain key areas of work at the interface of *in vitro* experimental and whole-system sampling-based approaches to microbial community ecology.

Another important direction for the future will be to explore how interactions between different species or strains can alter the dispersal behavior of co-habitating strains. In our work we found no evidence for *E. coli* or *E. faecalis* influencing the dispersal behavior of *P. aeruginosa*, but it is plausible that strains or species could influence others’ dispersal behavior through the secretion or adsorption of extracellular products. The degree to which dispersal is modulated by cross-species or cross-kingdom interactions and the ways in which these interactions influence species coexistence is an important frontier for future research that bears on fundamental microbial ecology. This topic also bears on infection scenarios in which the tendency of one species to disseminate from an initial site of a biofilm infection may depend on what other species are present as well.

---

**SI Figure S1:**
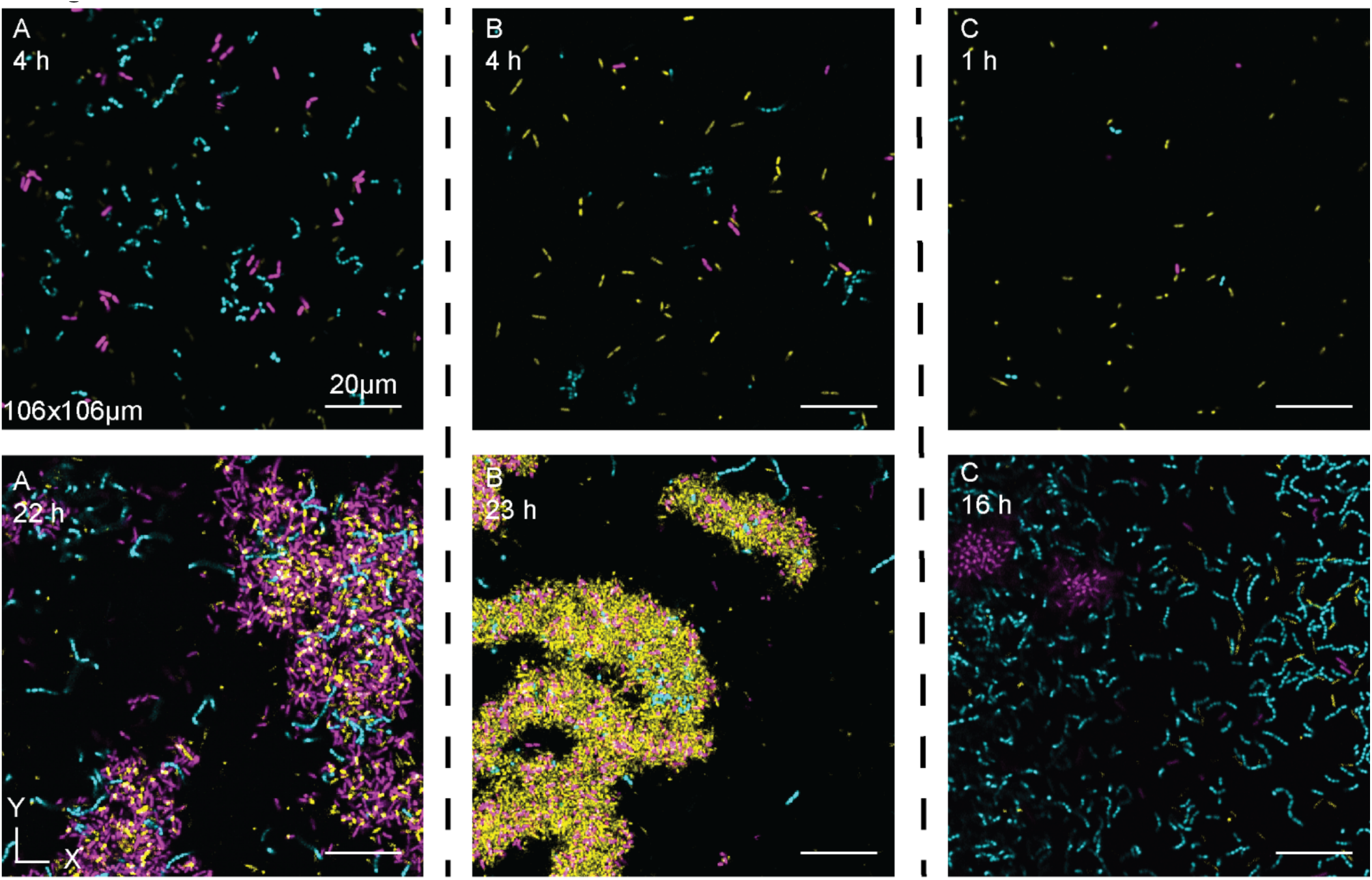
Representative images of variation in initial timepoints from three independent biofilm replicates. *P. aeruginosa* is shown in yellow, *E. coli* is shown in purple, and *E. faecalis* is shown in cyan. **(A)** Shows an *E. coli* dominated early time point. **(B)** Shows a *P. aeruginosa* dominated initial timepoint. **(C)** Shows an *E. faecalis* dominated early time point.

**SI Figure S2:**
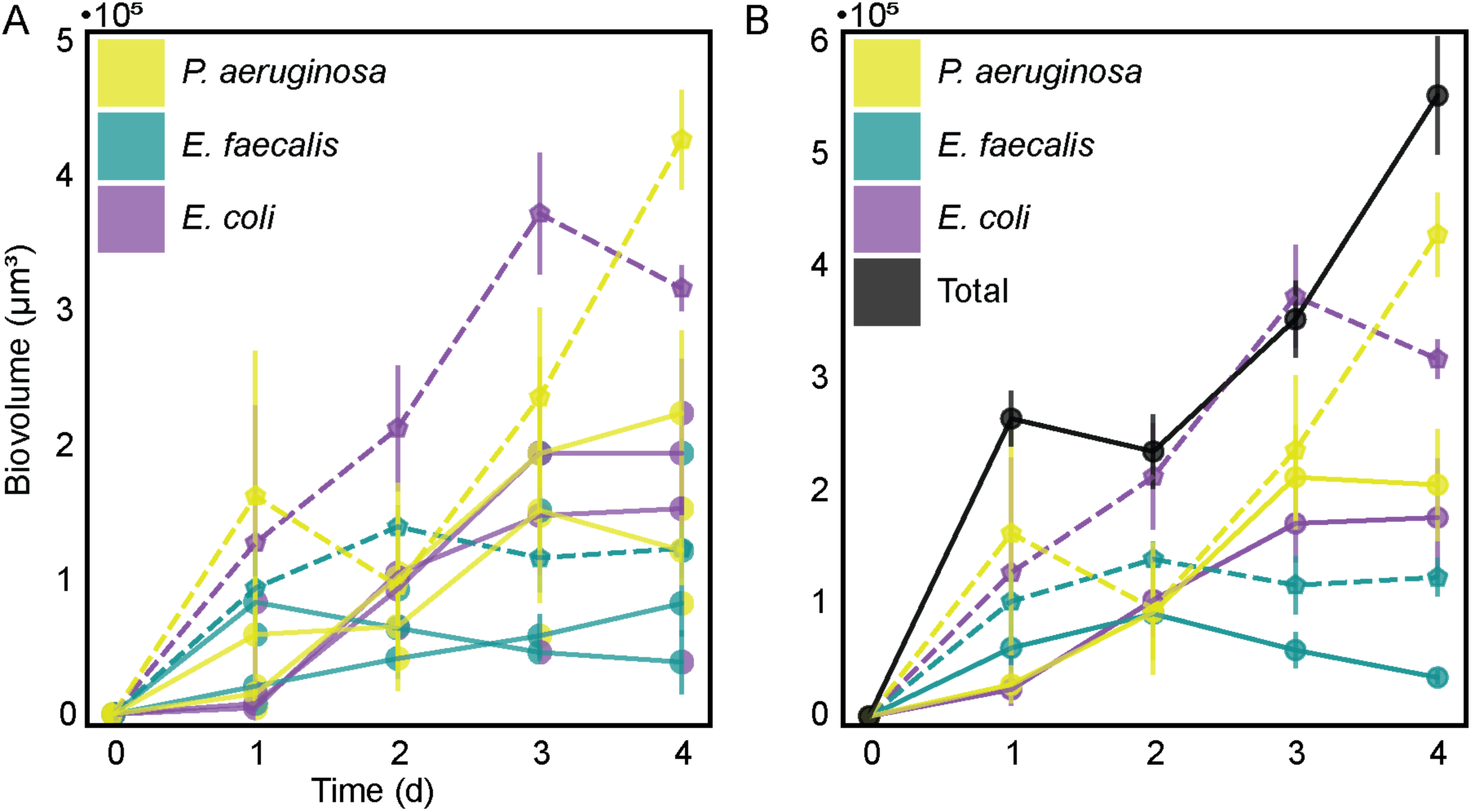
Competition in pairwise biofilm cocultures. **(A)** The biovolume for each species in co-culture (solid lines) plotted alongside monoculture biofilm control experiments (dashed lines) for each species across the first four days of growth (n=3-8). **(B)** The biovolume of each species in tri-culture (solid lines) plotted alongside monoculture biofilm control experiments (dashed lines) for each species across the first four days of growth. Dots represent the mean and error bars represent standard error (n=3-10). The color of the line represents the strain being plotted and the color of the dots represents the strains present in the culture. Solid colors represent monoculture, and split colors represent co-cultures. *P. aeruginosa* is shown in yellow, *E. coli* is shown in purple, and *E. faecalis* is shown in cyan. The sum of the three species biovolume when grown together as a community is shown in black.

**SI Figure S3:**
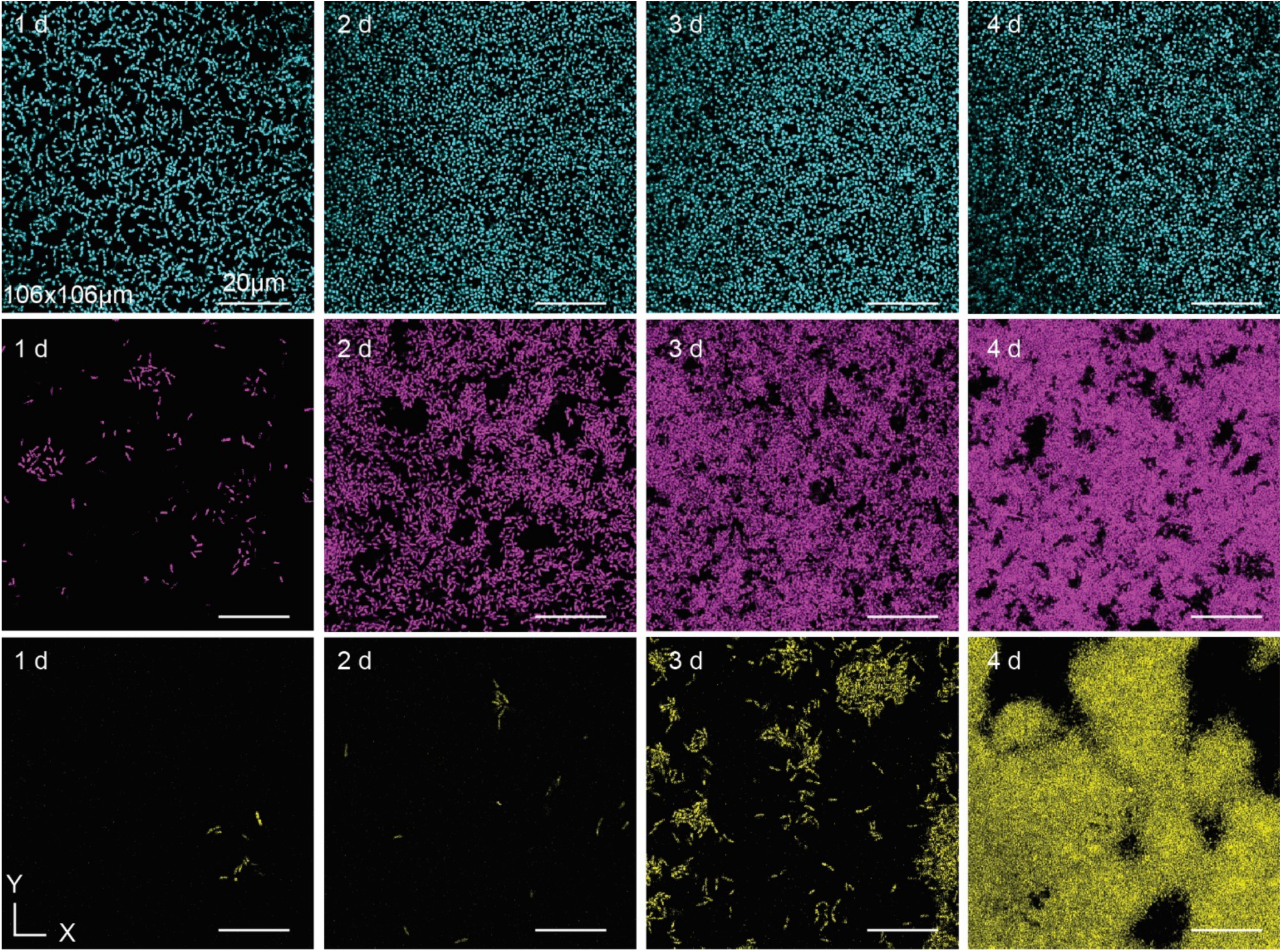
Representative images from the monoculture biofilm conditions. Top, *E. faecalis* monoculture biofilm (cyan). Middle, *E. coli* monoculture biofilm (purple). Bottom, *P. aeruginosa* monoculture biofilm (yellow).

**SI Figure S4:**
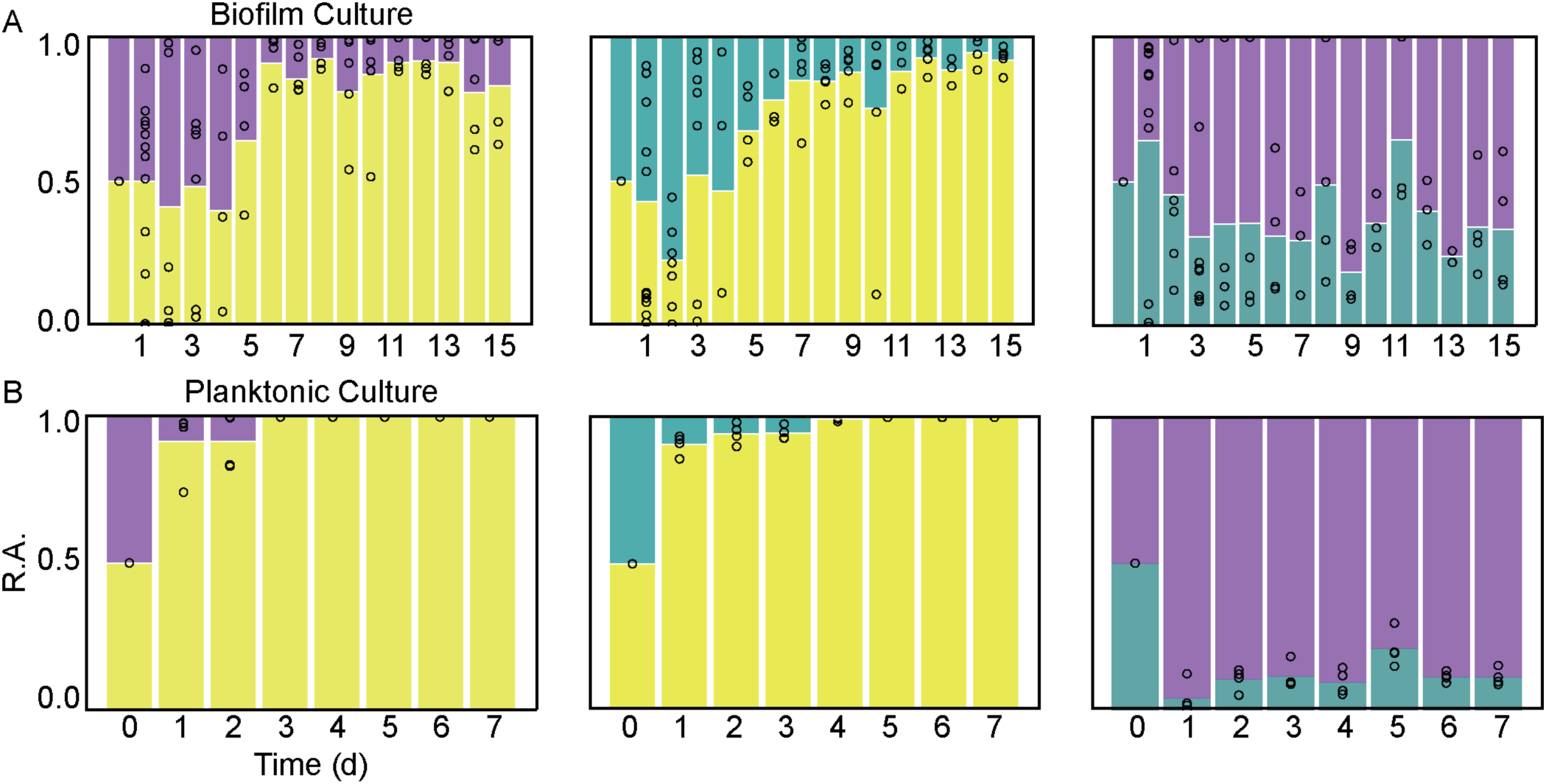
The species’ population dynamics in dual culture liquid competition are similar to the three-species coculture results. *P. aeruginosa* is shown in yellow, *E. coli* is shown in purple, and *E. faecalis* is shown in cyan. **(A)** Pairwise biofilm co-cultures of species relative abundance plotted against time (n=3-10). **(B)** Pairwise planktonic co-cultures of species relative abundance plotted against time (n=3-4).

**SI Figure S5:**
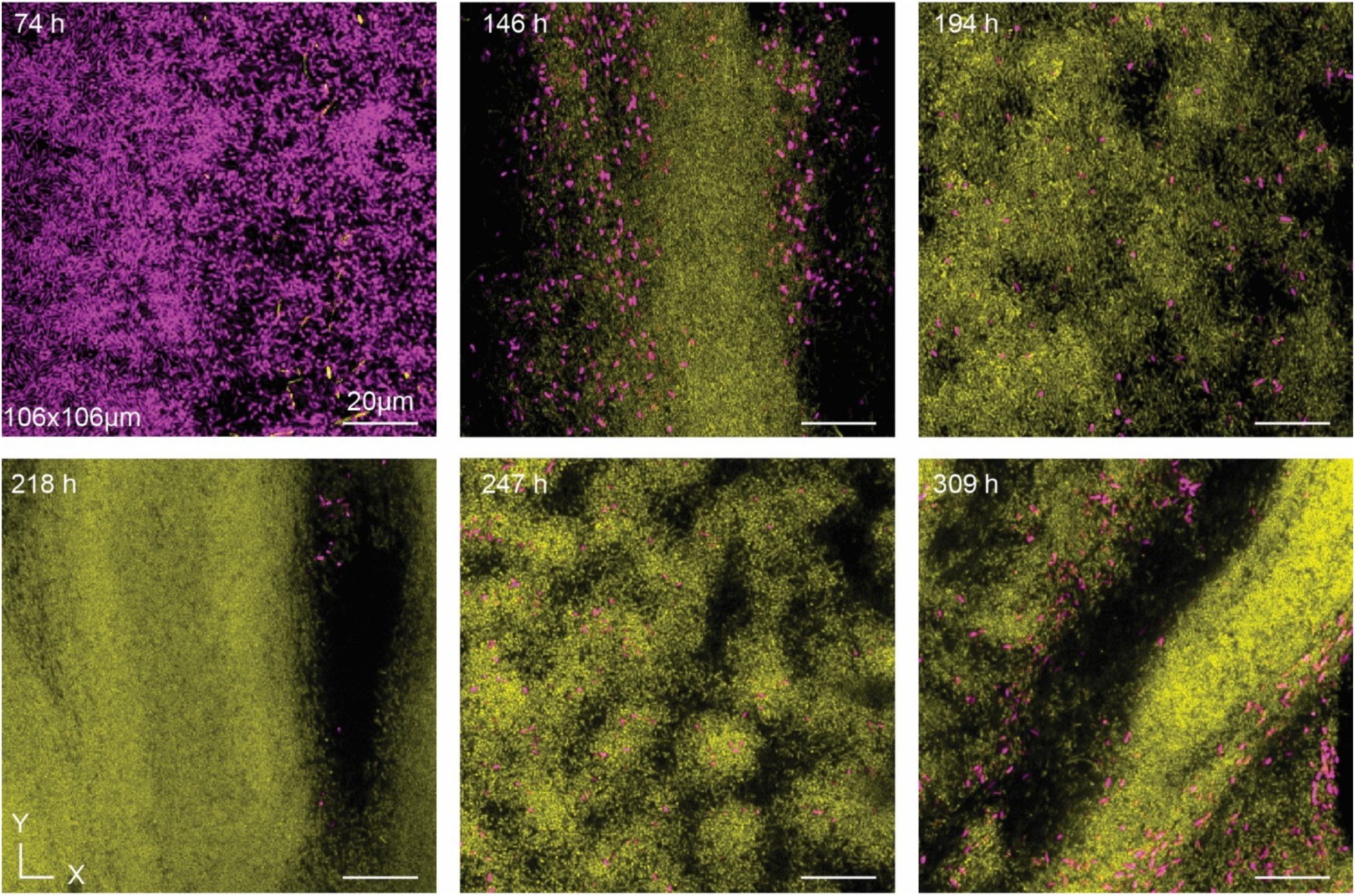
The *E. coli*-*P. aeruginosa* biofilm culture is not architecturally distinct from the triculture condition. Representative images of the *E. coli*-*P aeruginosa* biofilm time course show. *P. aeruginosa* is shown in yellow, *E. coli* is shown in purple, and *E. faecalis* is shown in cyan.

**SI Figure S6:**
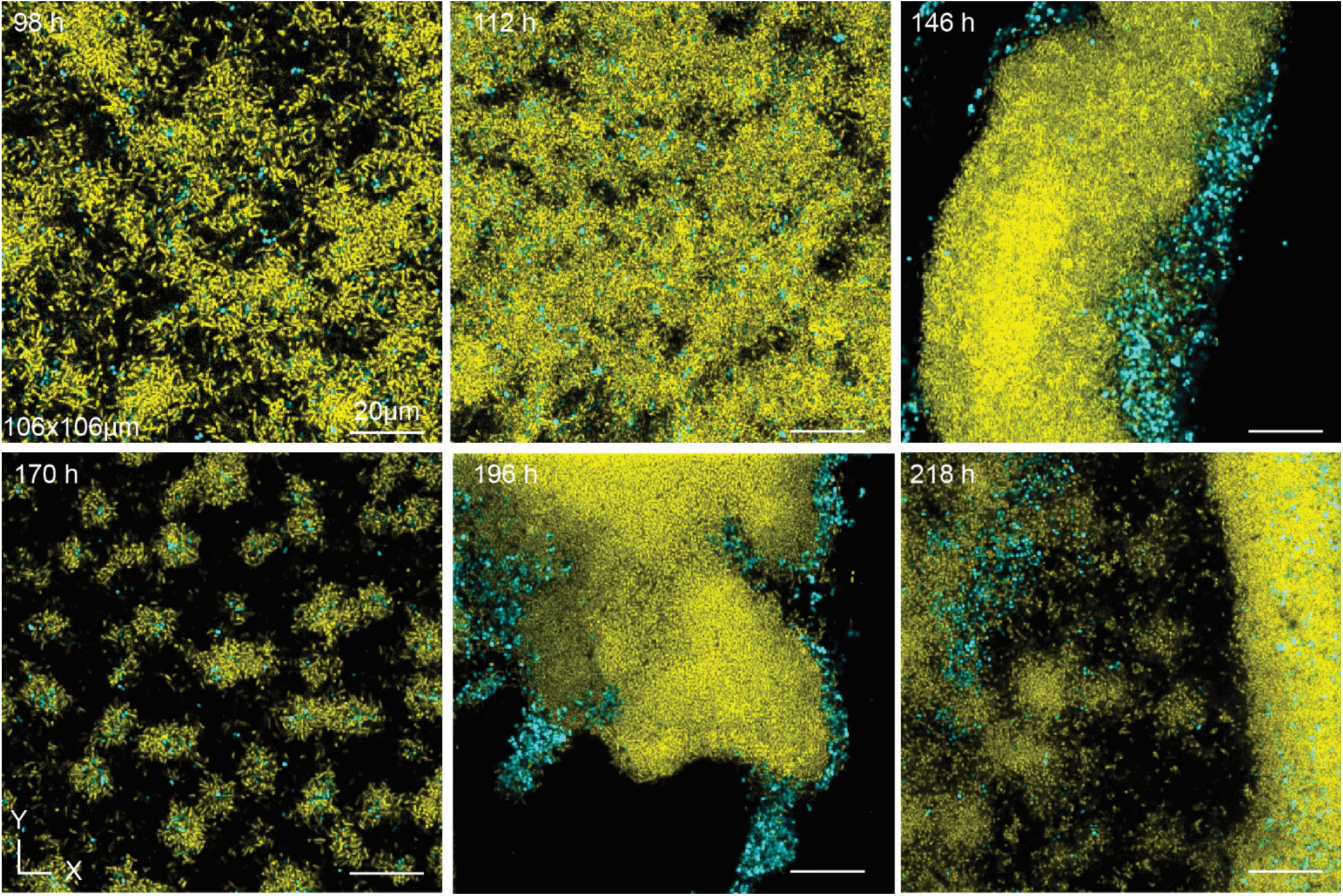
The *E. faecalis*-*P. aeruginosa* biofilm culture is not architecturally distinct from the triculture condition. Representative images of the *E. faecalis*-*P aeruginosa* biofilm time course. *P. aeruginosa* is shown in yellow, *E. coli* is shown in purple, and *E. faecalis* is shown in cyan.

**SI Figure S7:**
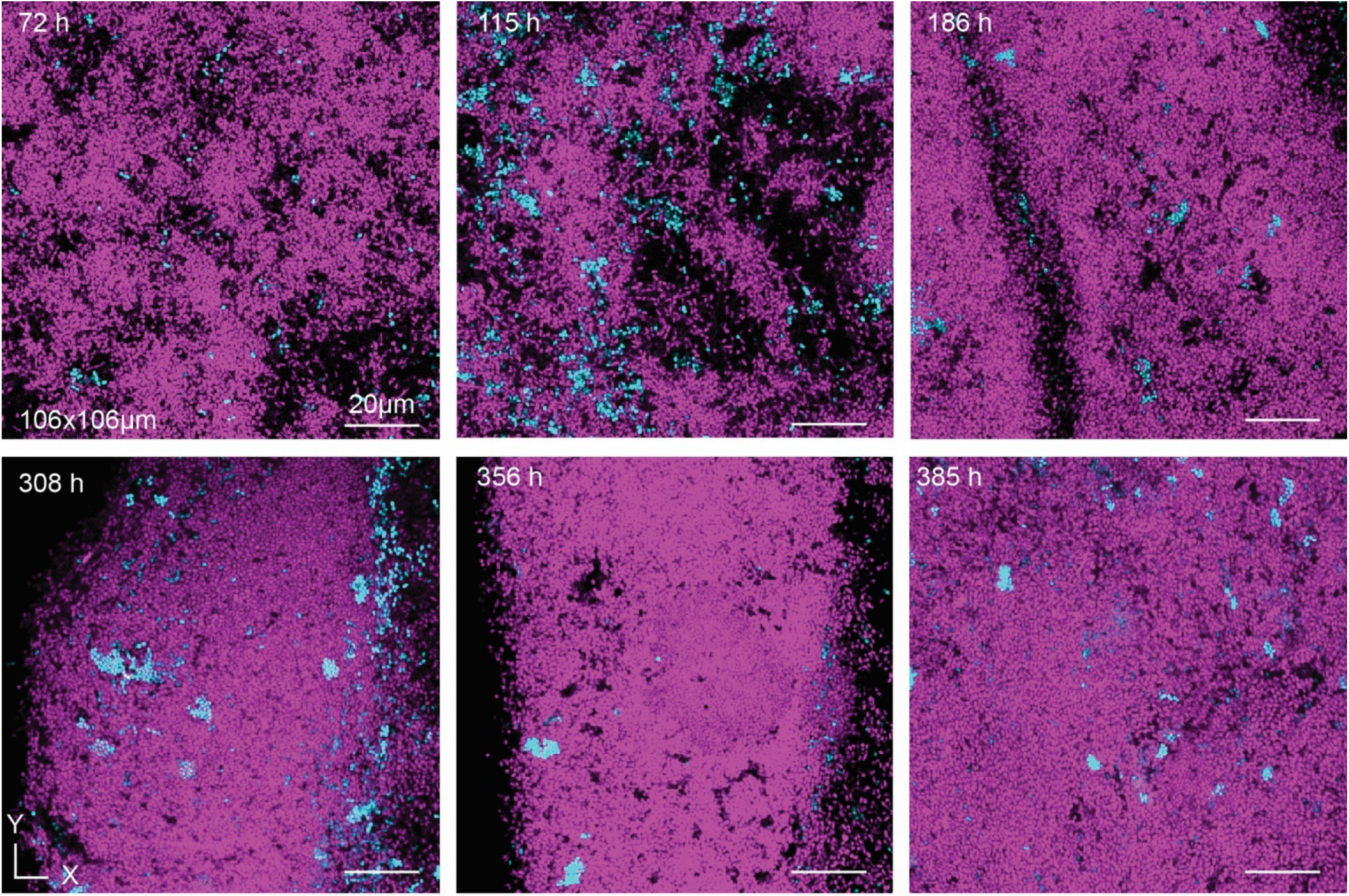
The *E. coli-E. faecalis* biofilm is dominated by *E. coli* and has a relatively stable architecture in comparison to biofilm cultures containing *P. aeruginosa*. Representative images of the *E. coli-E. faecalis* biofilm time course. *P. aeruginosa* is shown in yellow, *E. coli* is shown in purple, and *E. faecalis* is shown in cyan.

**SI Figure S8:**
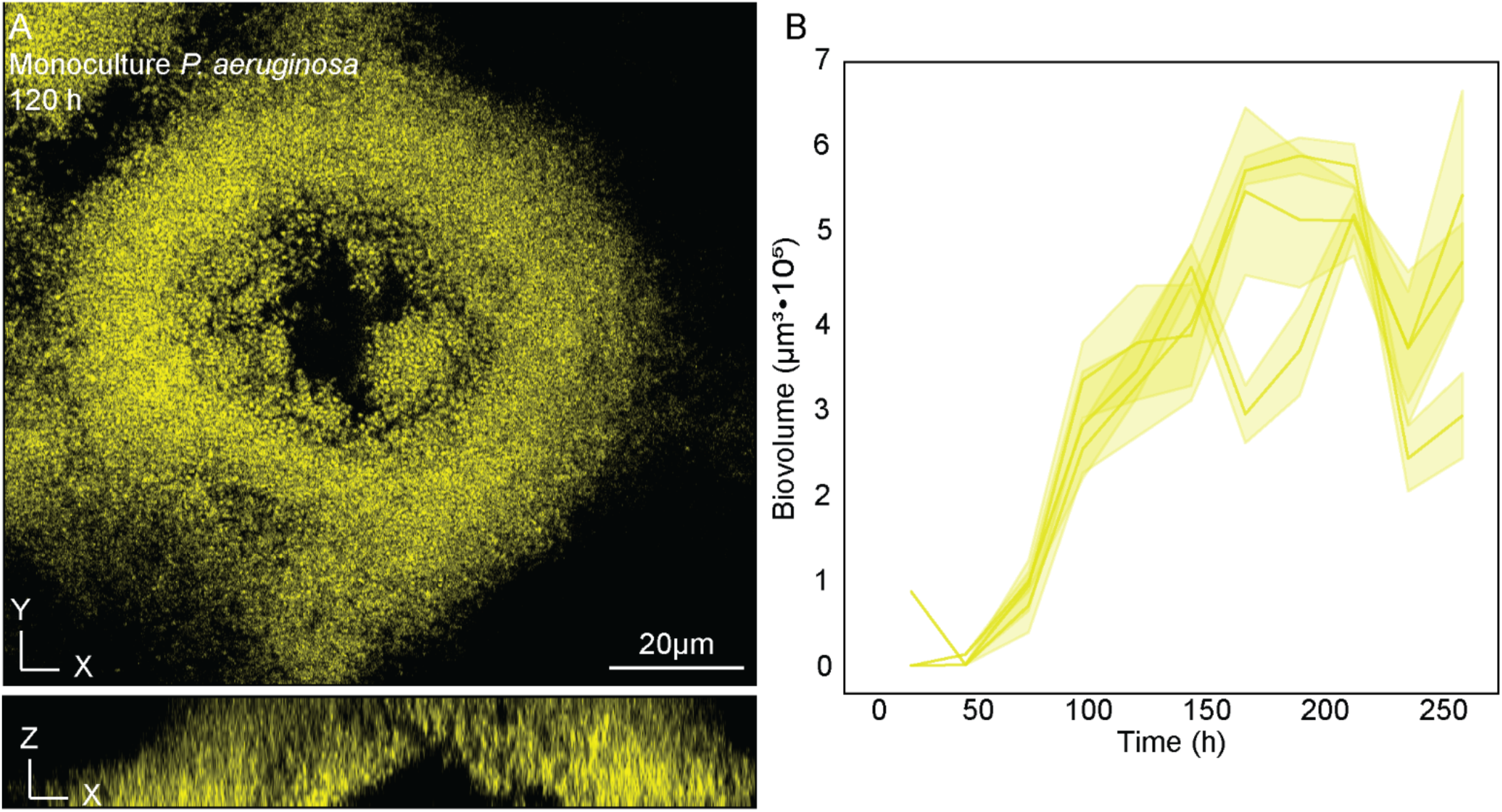
Monoculture *P. aeruginosa* (yellow) disperses from the microfluidic chamber. **(A)** Representative image of a hollowed *P. aeruginosa* colony, characteristic of *P. aeruginosa* biofilm dispersal. **(B)** Traces of *P. aeruginosa* monoculture biofilms. Solid lines represent the mean and shaded regions represent one standard deviation above and below the mean (n = 3).

**SI Figure S9:**
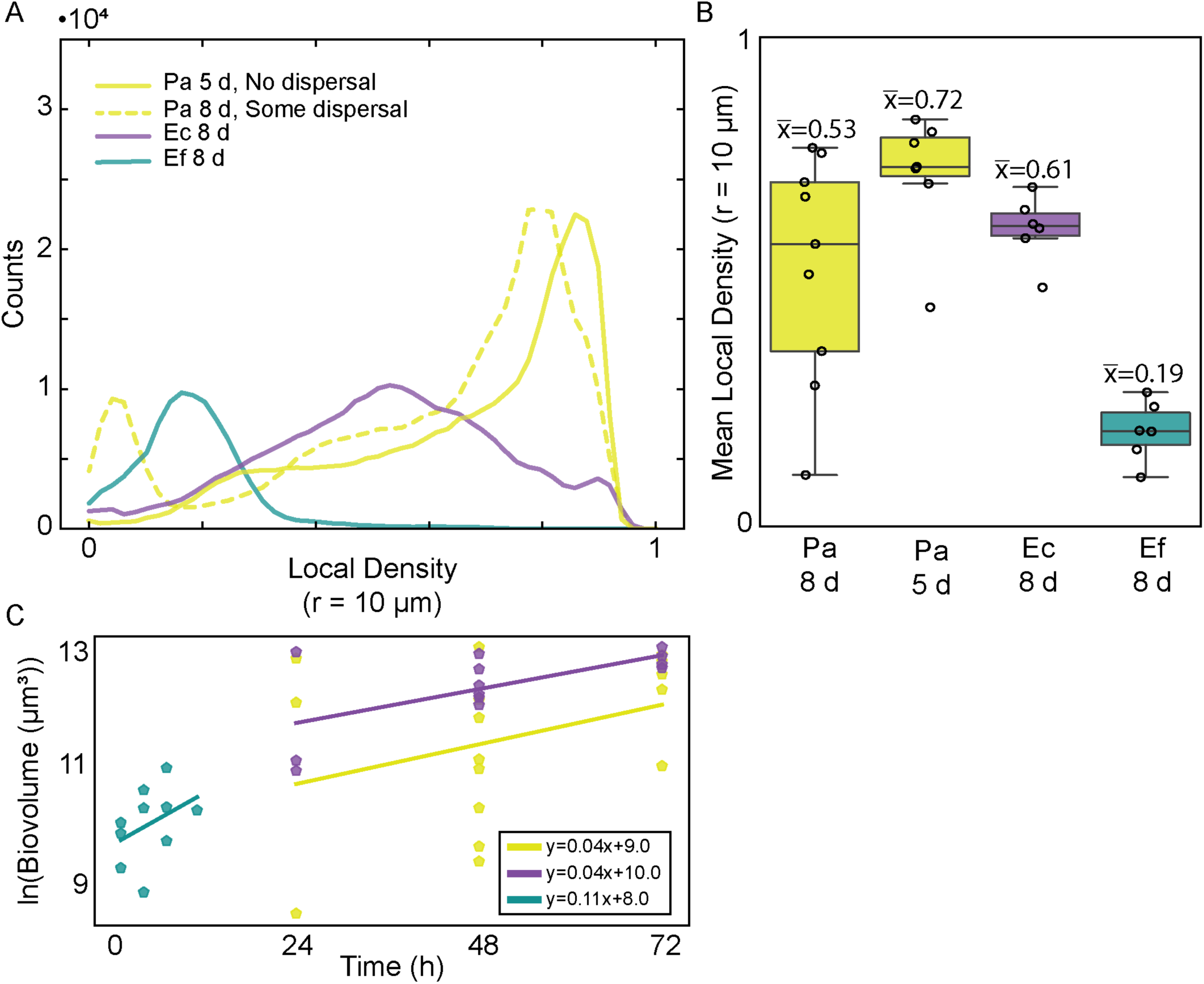
The three species achieve a range of growth rates and mean local density when grown in monoculture. **(A)** Histograms of mean local density (radius = 10 µm) for *E. coli* (purple), *P. aeruginosa* (yellow), and *E. faecalis* (cyan) (n=3-4). **(B)** Corresponding box and whisker plots of mean local density (radius = 10 µm) for *E. coli*, *P. aeruginosa*, and *E. faecalis* (n=3-4). **(C)** The maximal growth rate was determined by taking the slope of the natural log of the biovolume of monoculture biofilms during initial timepoints (n= 3-8).

**SI Figure S10:**
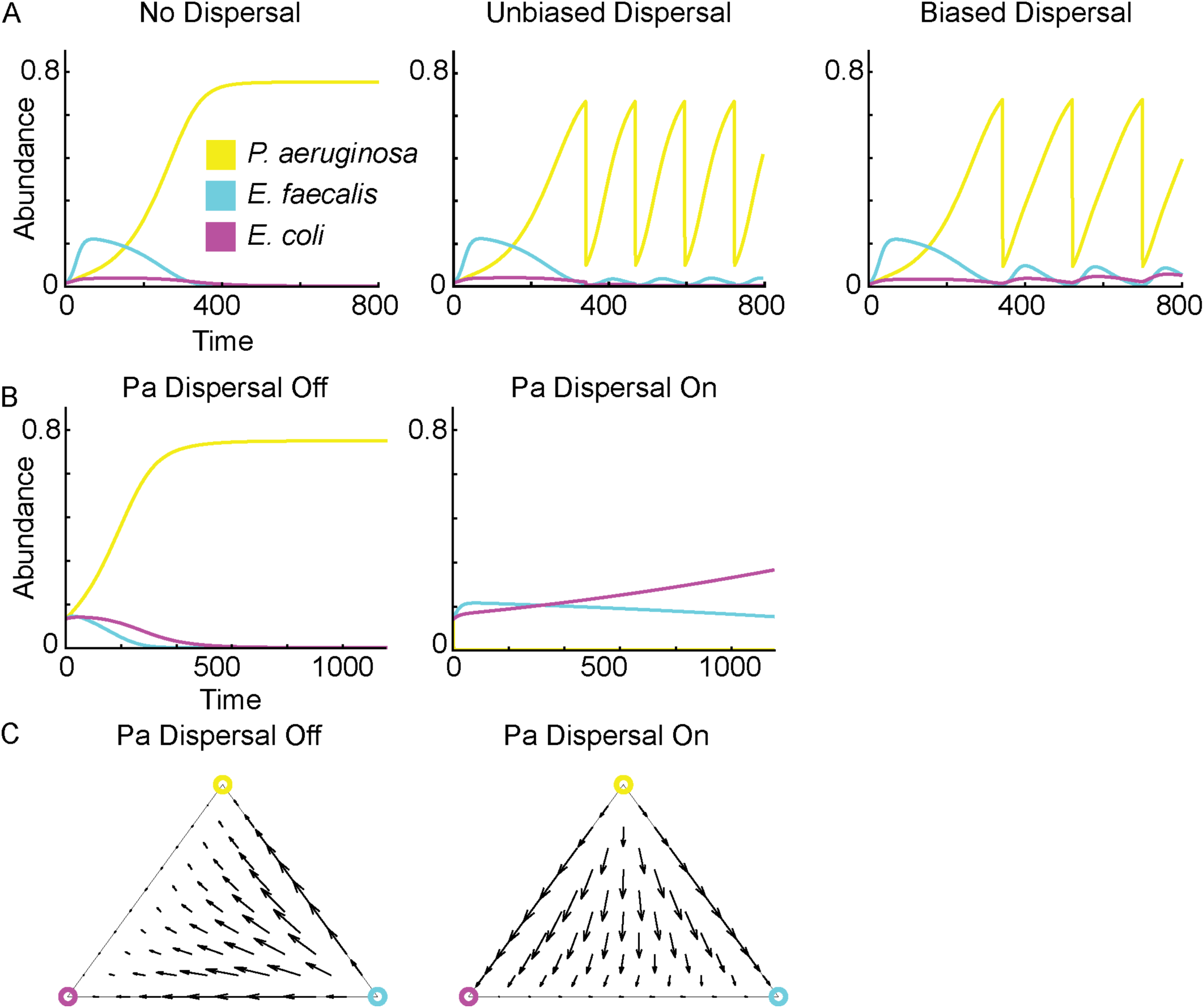
A mean-field model supports biased dispersal of the dominant competitor as a mechanism of coexistence. **(A)** Biased dispersal of *P. aeruginosa* results in coexistence of the three species. **(B)** Trajectories of the biased dispersal case when dispersal is turned on and when dispersal is turned off. **(C)** Ternary vector-plots of the biased dispersal case when *P. aeruginosa* dispersal is turned on and when *P. aeruginosa* dispersal is turned off. **(D)** Phase diagram of the biased dispersal model plotting *P. aeruginosa*’s carrying capacity, K_Pa_, against dispersal ratio (*δ_j_*/*δ_Pa_*), ranging from the Biased Dispersal case (dispersal ratio of 0) to the Unbiased Dispersal case (dispersal ratio of 1) as the carrying capacity of *P. aeruginosa* (*K_Pa_*) is varied from 0.05 to 3.0. The color map represents *P. aeruginosa*’s relative abundance after 4000 simulation time steps.

**SI Figure S11:**
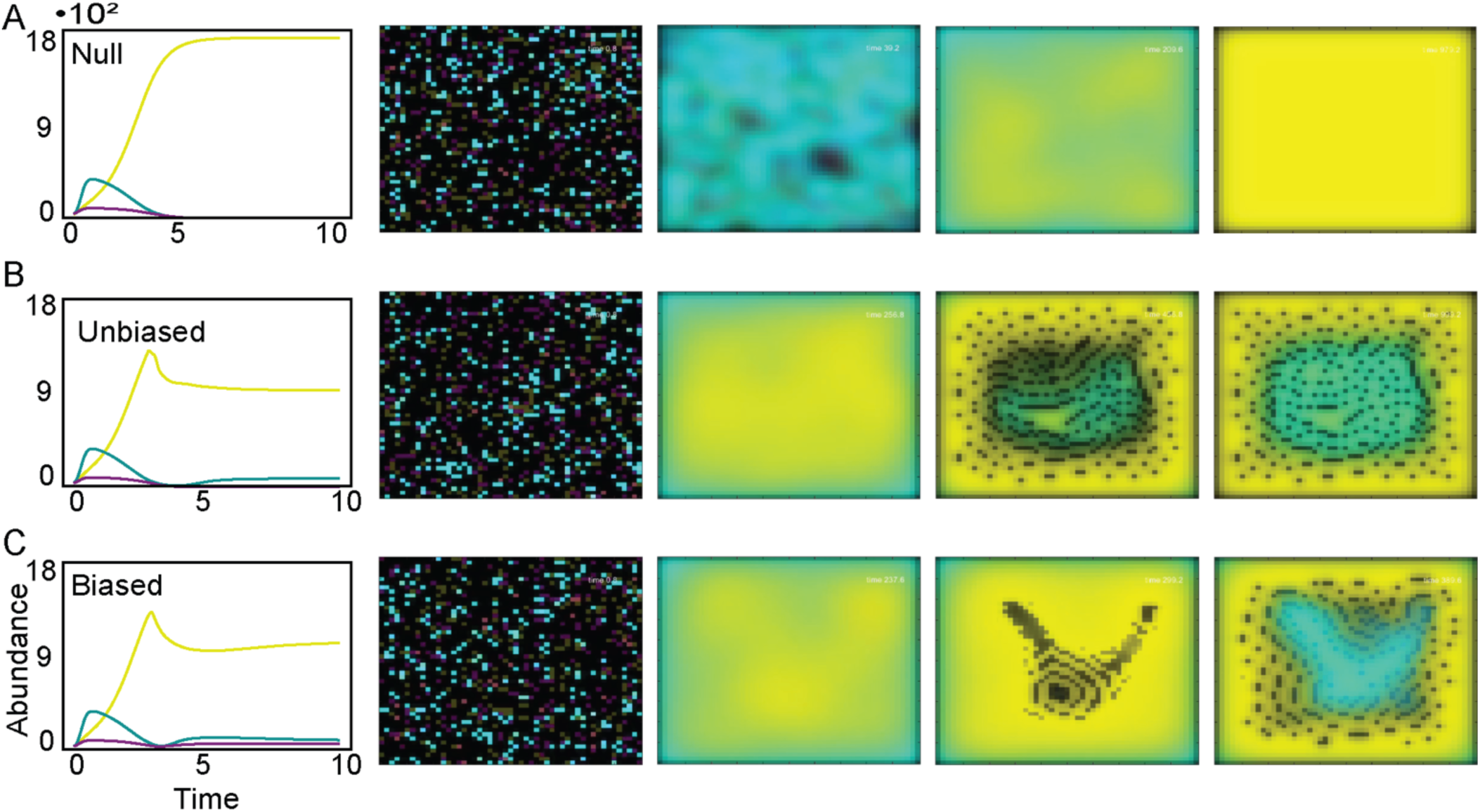
Coexistence in the deterministic-spatial model is dependent on dispersal bias. **(A)** No dispersal occurs. **(B)** All three species disperse equally. **(C)** Only *P. aeruginosa* disperses.

**SI Figure S12:**
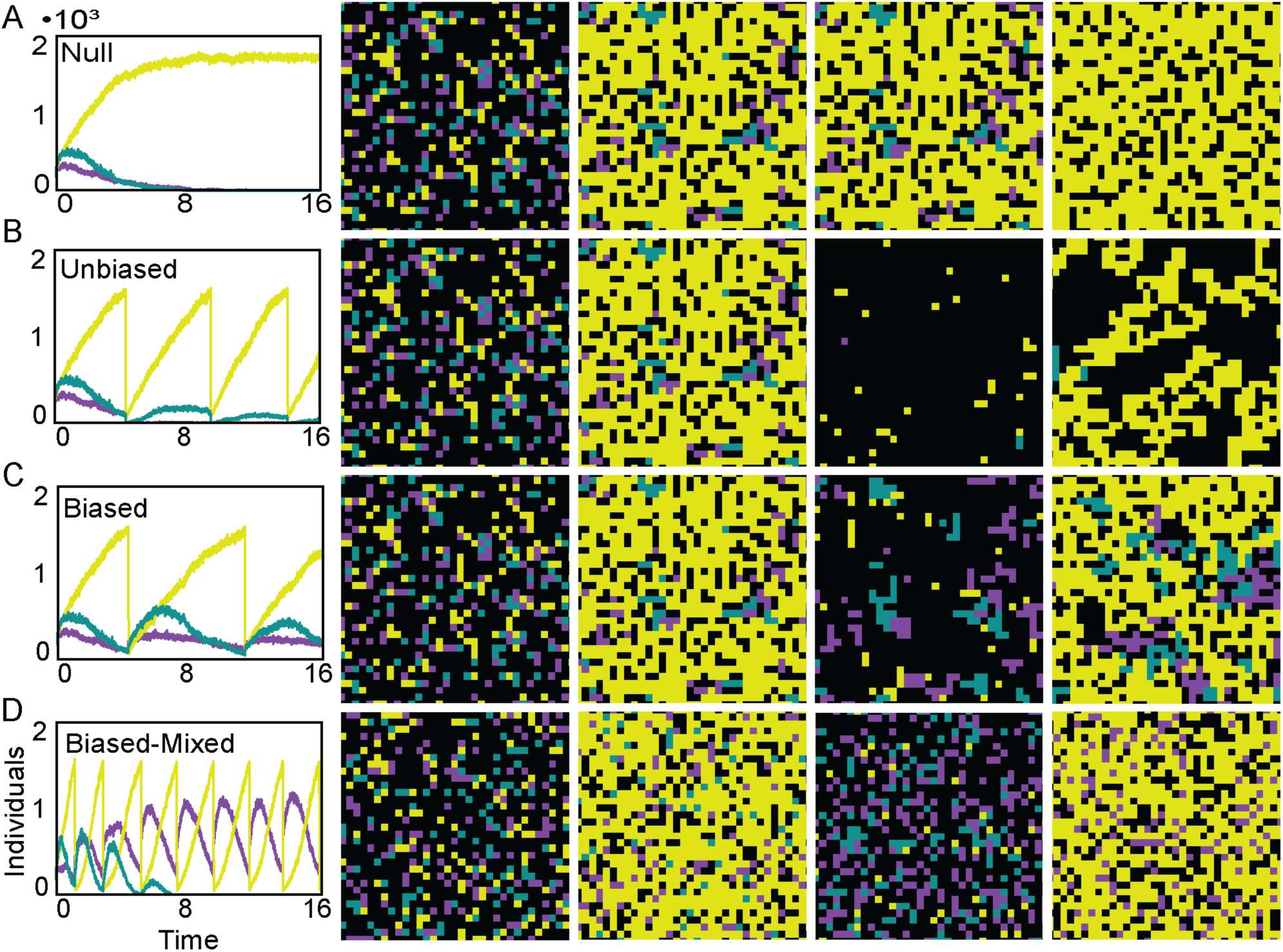
Coexistence in the stochastic-spatial model is dependent on dispersal bias and spatial arrangement. **(A)** No dispersal occurs. **(B)** All three species disperse equally. **(C)** Only *P. aeruginosa* disperses. **(D)** Random mixing in the biased dispersal case leads to a loss of *E. faecalis*.

**SI Figure S13:**
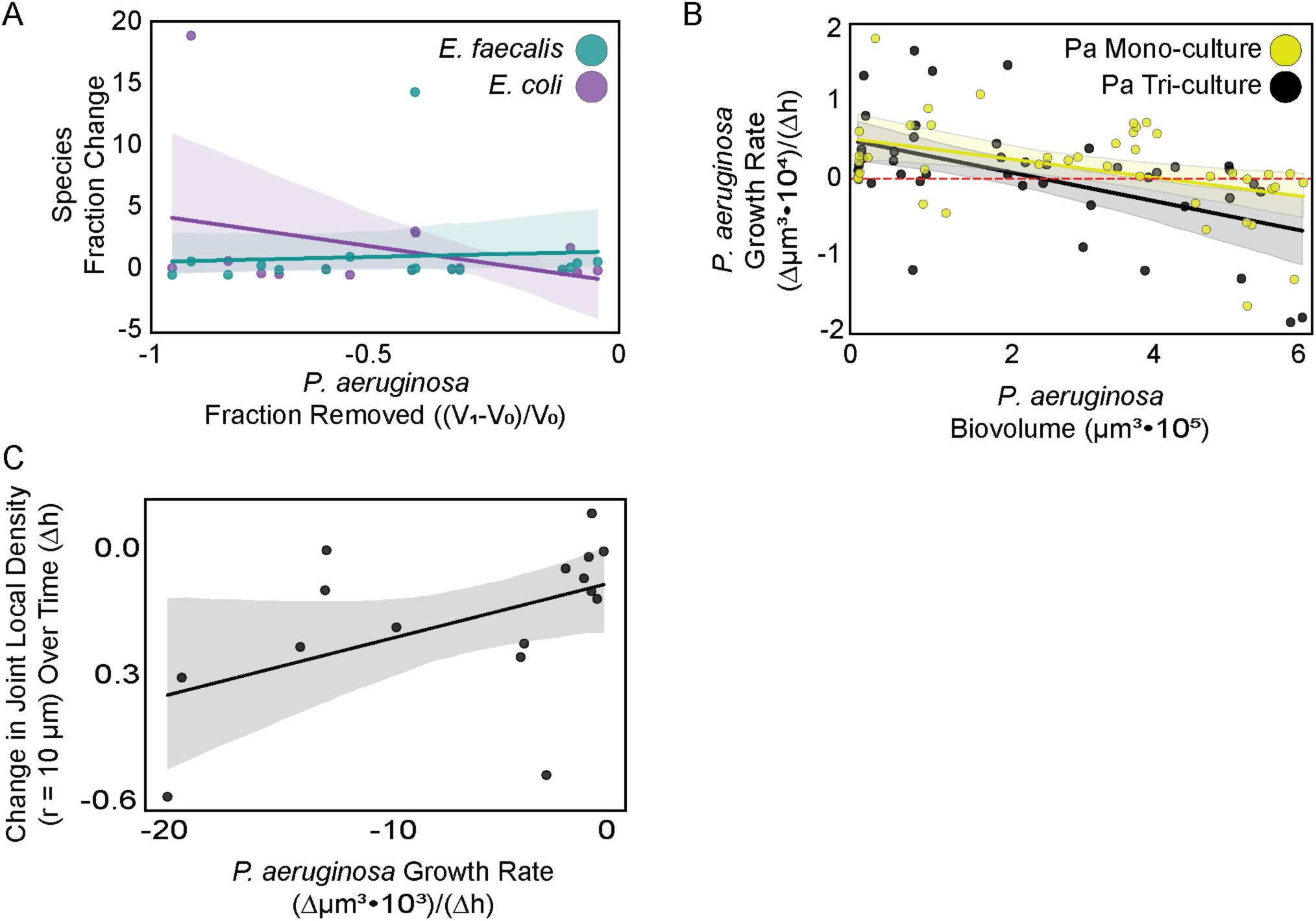
*P. aeruginosa* drives dispersal events, altering biofilm architecture. **(A)** *P. aeruginosa* dispersal is biased in the sense that there is no correlation between the fraction of *P. aeruginosa* biovolume removed and the fraction of *E. coli* (r^2^=0.09, p=0.1911, data from n = 3 independent experiments) or *E. faecalis* (r^2^=0.01, p=0.7930, data from n = 3 independent experiments) biovolume removed. **(B)** Monoculture *P. aeruginosa* disperses in a population density-dependent manner that is not significantly different than when in the three-species community. 95% confidence interval (shaded region) of linear regression fits overlap between monoculture and coculture dataset (mono-culture linear regression r^2^=0.16, p=0.0032, tri-culture linear regression, r^2^ = 0.25, p = 0.0002, data from n = 3 independent experiments). **(C)** *P. aeruginosa* dispersal events correlate moderately with changes in total biofilm density (r^2^=0.25, p=0.0489, data from n = 3 independent experiments). Solid lines represent line fits and shaded regions represent a 95% confidence interval.

**SI Figure S14:**
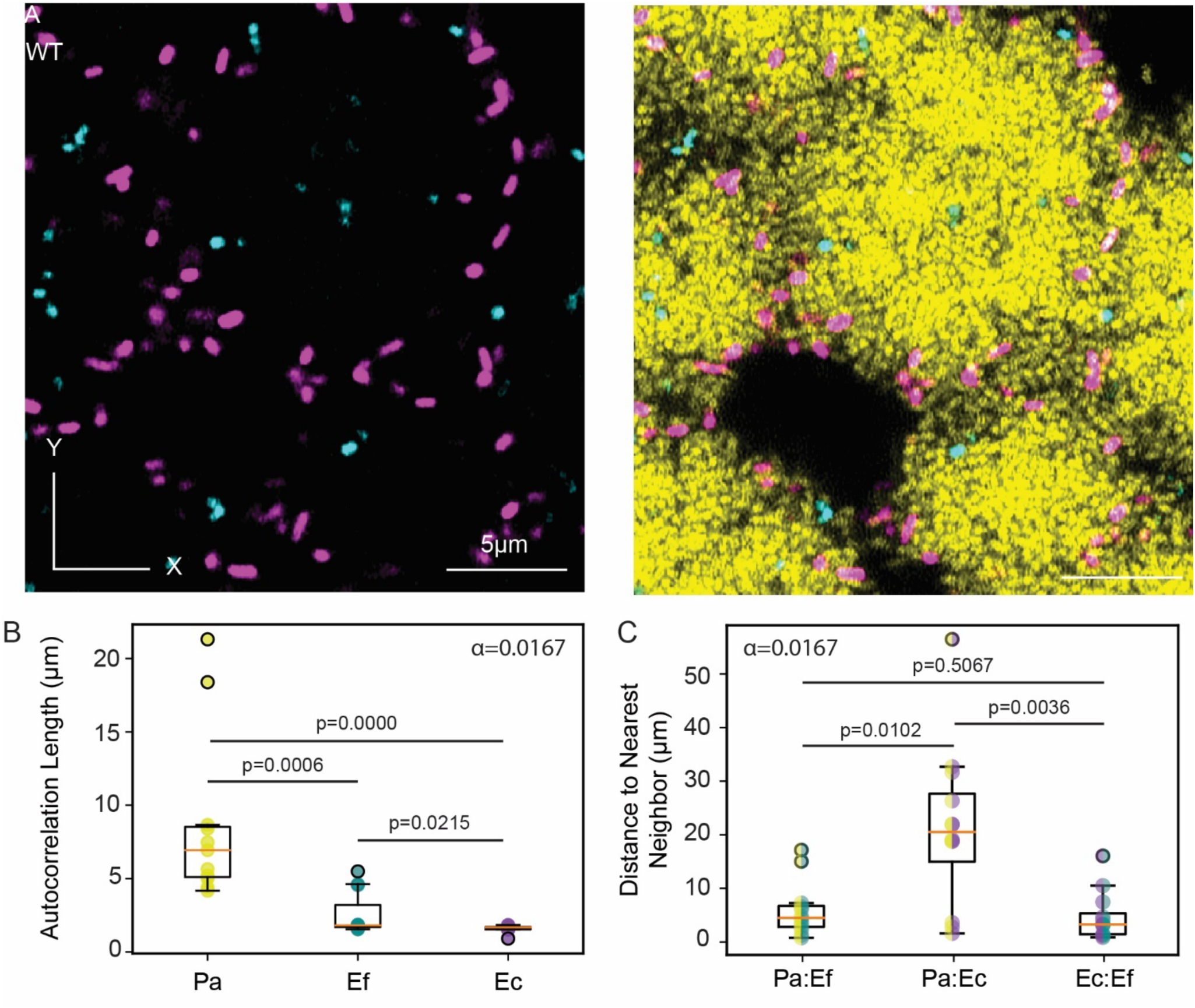
Spatial localization of the three species relative to themselves and one-another. **(A)** Representative image of the community biofilm. *P. aeruginosa* is shown in yellow, *E. coli* is shown in purple, and *E. faecalis* is shown in cyan. **(B)** Autocorrelation length of each of the three species in the community at 15 d (Mann-Whitney U tests, Bonferroni corrected α, n = 11 locations from 3 biofilm chambers). **(C)** Distance to nearest neighbor in the community at 15 d (Mann-Whitney U tests, Bonferroni corrected α, n = 11 locations from 3 chambers).

**SI Figure S15:**
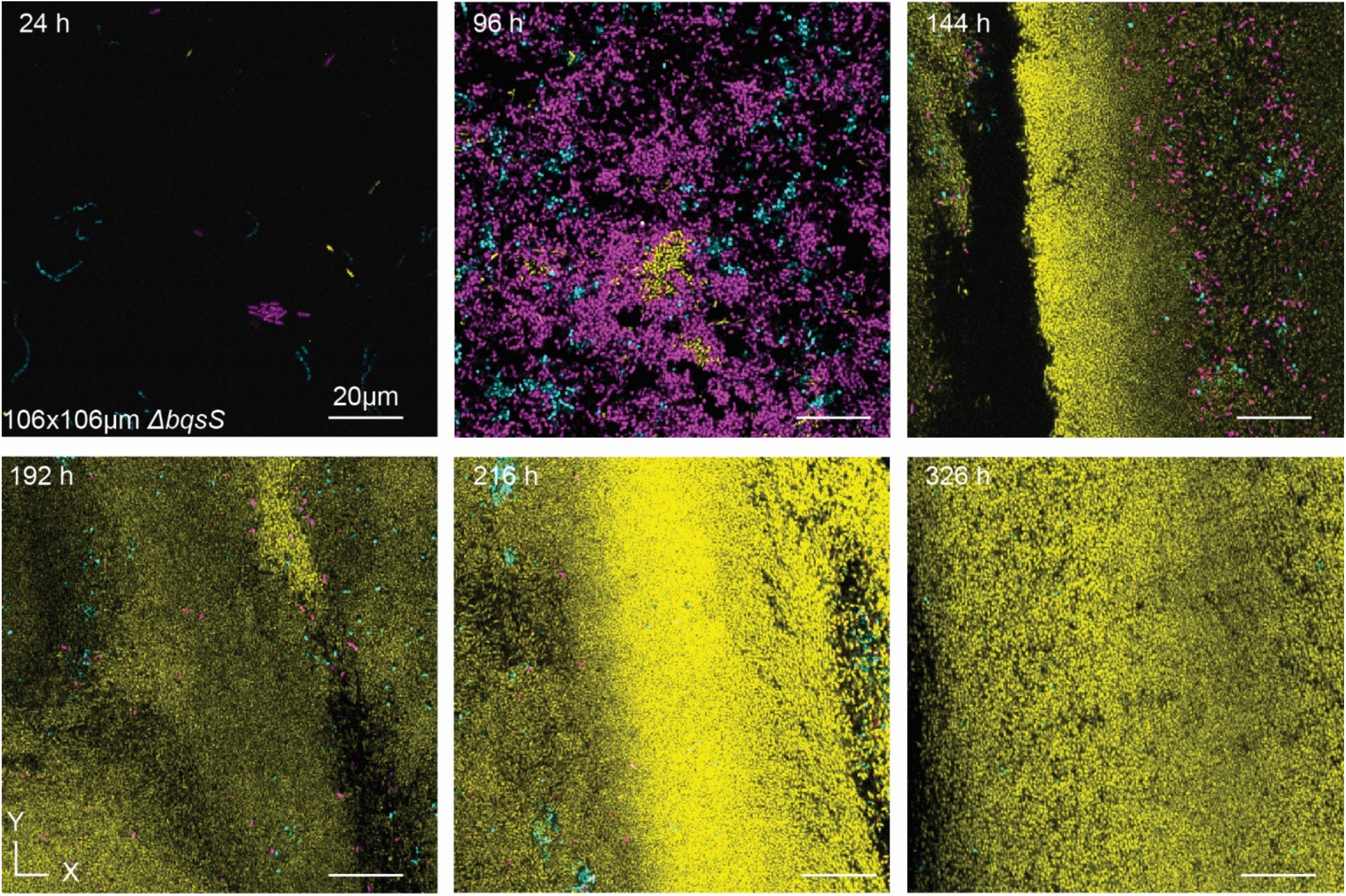
The *P. aeruginosa ΔbqsS* biofilm community does not undergo mass dispersal events. Representative images of the *P. aeruginosa ΔbqsS* biofilm community time course showing consistent biofilm architecture and decreased abundance of *E. coli* and *E. faecalis* relative to *P. aeruginosa* WT biofilm communities. *P. aeruginosa* is shown in yellow, *E. coli* is shown in purple, and *E. faecalis* is shown in cyan.

**SI Figure S16:**
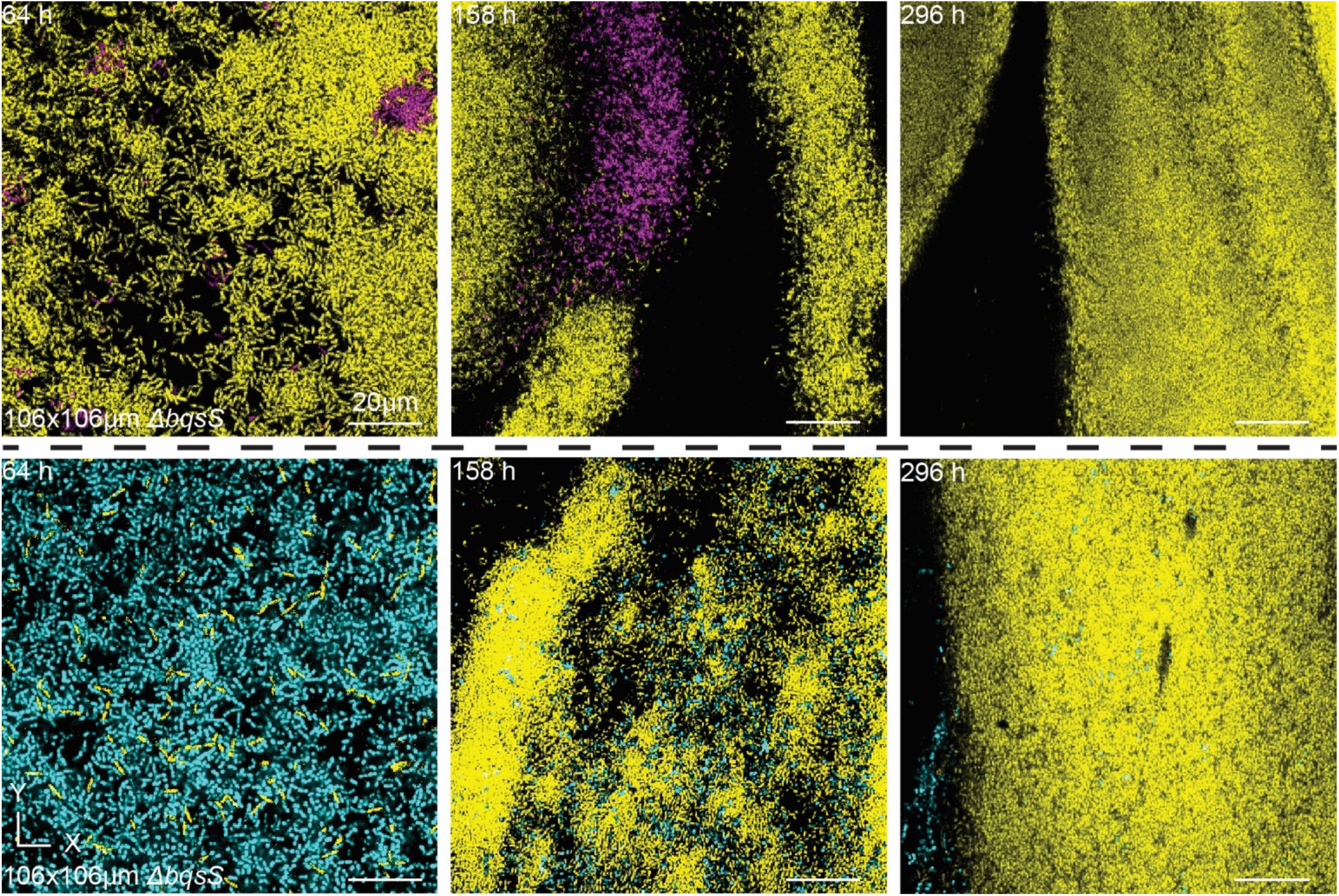
Pairwise biofilm cocultures with the *P. aeruginosa ΔbqsS* mutant do not undergo mass dispersal events. Representative images of the *P. aeruginosa ΔbqsS* pairwise biofilm cultures showing consistent biofilm architecture and decreased abundance of *E. coli* and *E. faecalis* relative to *P. aeruginosa* WT biofilm cocultures. *P. aeruginosa* is shown in yellow, *E. coli* is shown in purple, and *E. faecalis* is shown in cyan.

**SI Figure S17:**
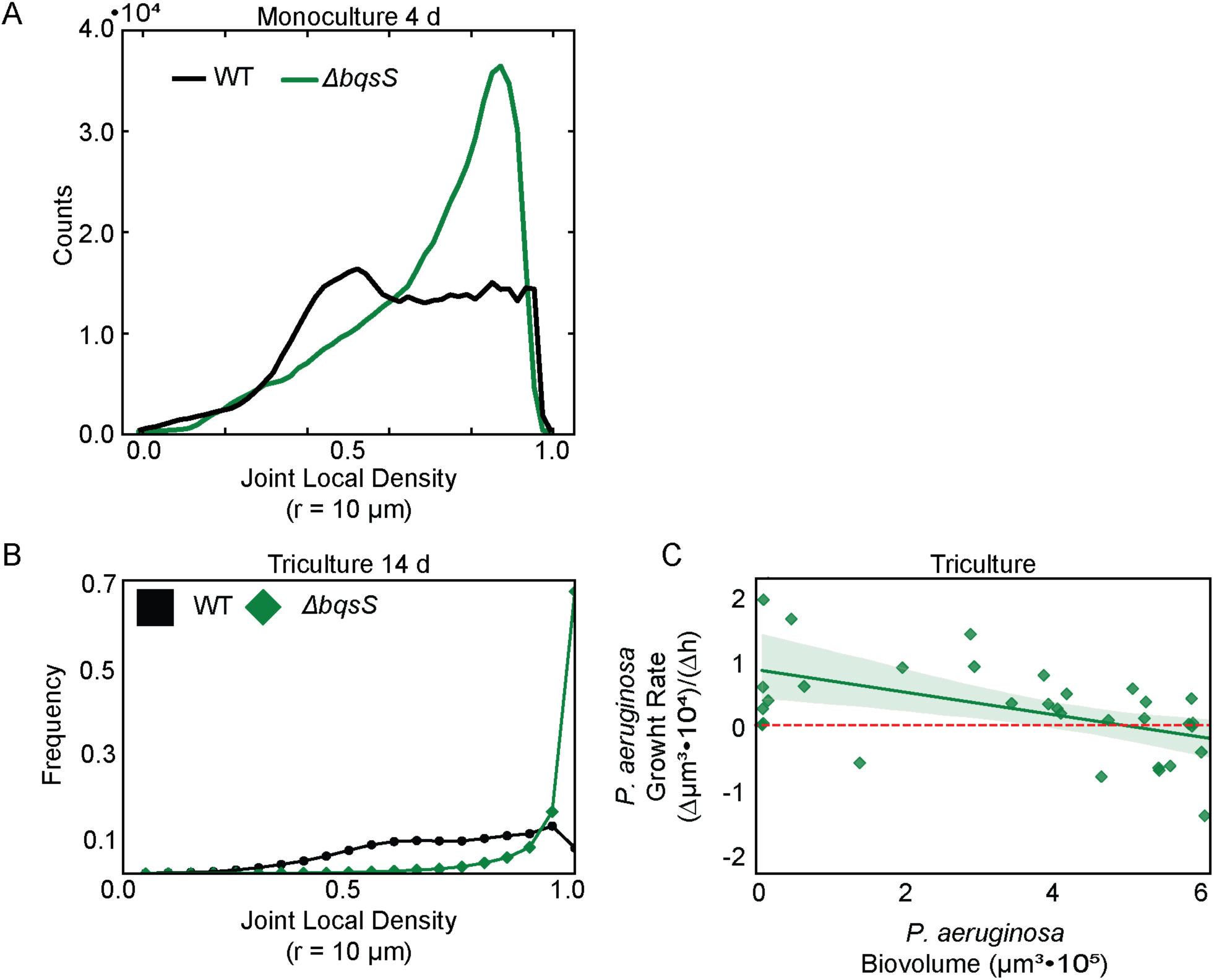
*P. aeruginosa ΔbqsS* has a shifted density profile and does not disperse in the same density-dependent manner as WT *P. aeruginosa*. **(A)** Histograms of local population density (r = 10 μm) of *ΔbqsS* and WT *P. aeruginosa* monoculture biofilms (n = 3). **(B)** Histograms of joint local population density (r = 10 μm) of *ΔbqsS* and WT *P. aeruginosa* biofilm communities (n = 4). **(C)** *P. aeruginosa ΔbqsS* biovolume does correlate with its change in biovolume at the following timestep (r^2^ = 0.25, p = 0.0013, n = 34 timesteps), but crosses 0 (dashed, red line) at a higher value (n = 3). The solid line represent the line fit and the shaded region represents a 95% confidence interval.

**SI Figure S18:**
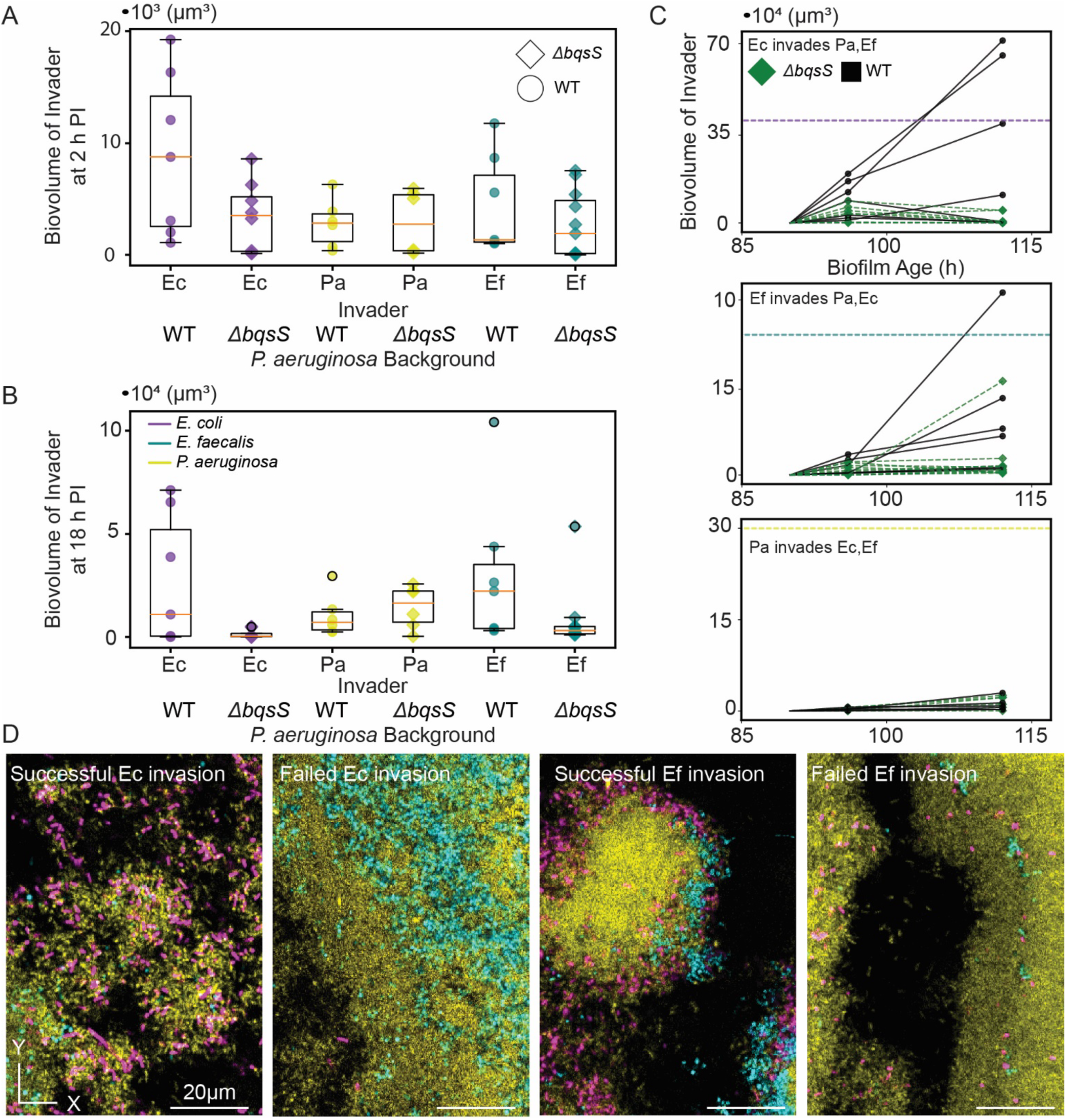
*P. aeruginosa* is conditionally susceptible to invasion by *E. coli* and *E. faecalis*. **(A)** Box and whisker plots of the biovolume of the invading strain 2 h post invasion (n = 6-10). **(B)** Box and whisker plots of the biovolume of the invading strain 18 h post invasion (n = 6-10). **(C)** Trajectories of the invading strain biovolume with the average biovolume from the three-species biofilm competition experiments overlayed as a dashed line (n = 6-10). **(D)** Representative images of successful and failed invasions by *E. coli* and *E. faecalis* into biofilm cocultures containing WT *P. aeruginosa*.

**SI Table S1:**
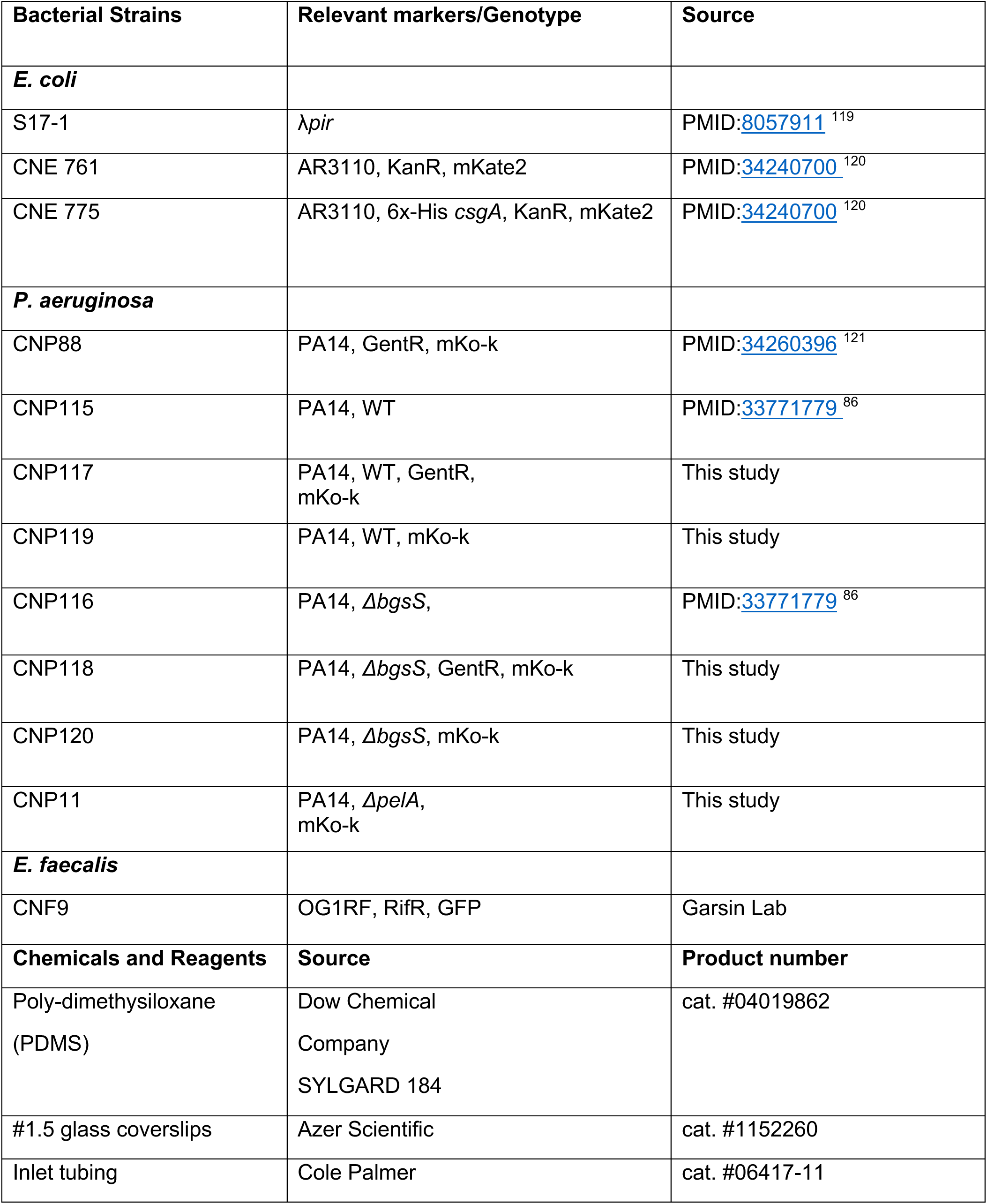
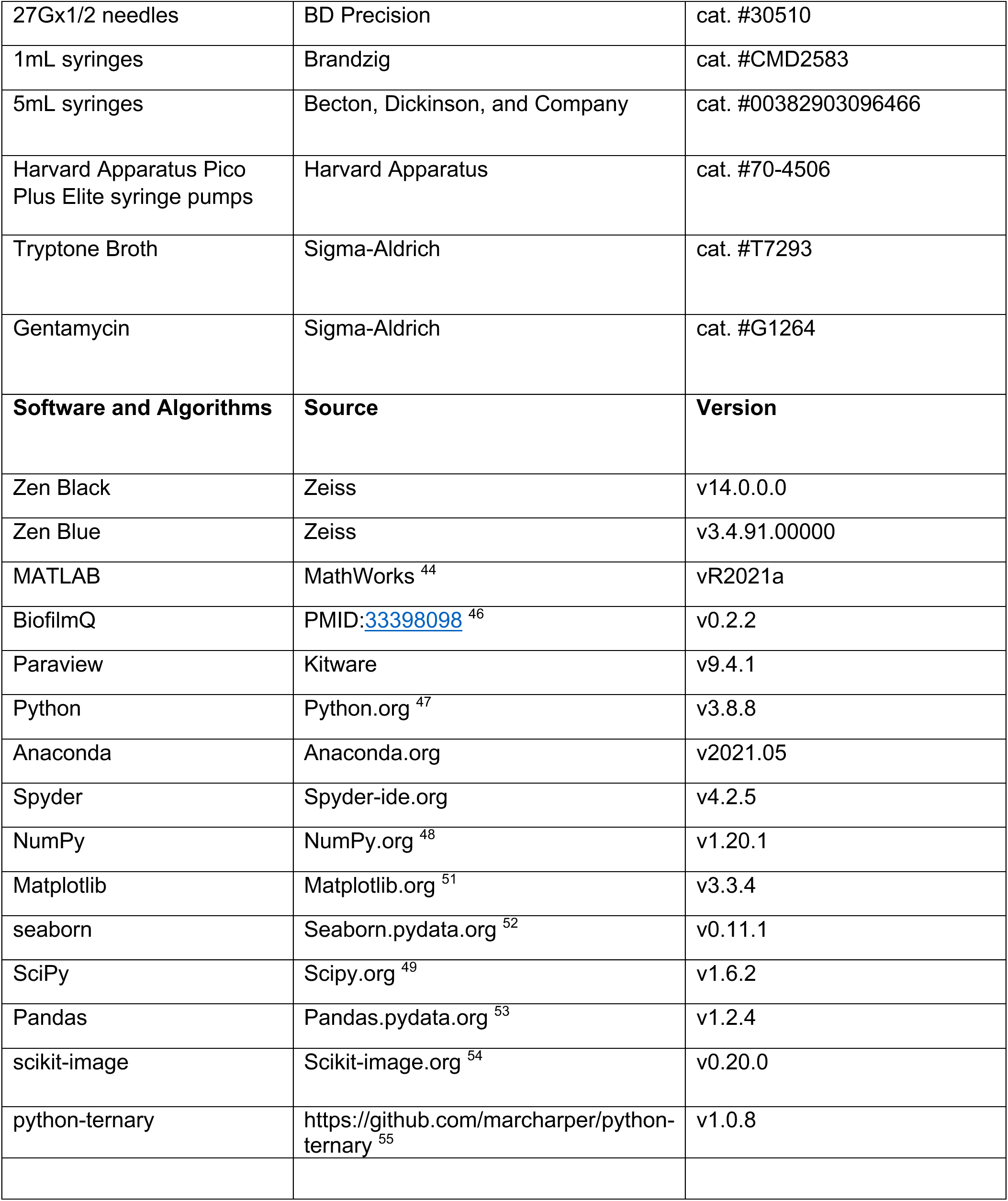
List of strains, materials, and software.

**SI Table S2:** List of model parameters.

| Parameter | Symbol | Value |
| --- | --- | --- |
| *P. aeruginosa* maximal growth rate | $r_{Pa}$ | 0.042 |
| *E. faecalis* maximal growth rate | $r_{Ef}$ | 0.109 |
| *E. coli* maximal growth rate | $r_{Ec}$ | 0.036 |
| *P. aeruginosa* carrying capacity | $K_{Pa}$ | 0.72 |
| *E. faecalis* carrying capacity | $K_{Ef}$ | 0.19 |
| *E. coli* carrying capacity | $K_{Ec}$ | 0.61 |
| Dispersal rate | $\delta_i$ | 0.2 |
| Dispersal trigger | $N_{down}$ | $0.9 * K_{pa}$ |
| End of dispersal trigger | $N_{up}$ | $0.1 * K_{pa}$ |

## SI Text

### Mathematical models

Our models build from the classical Lotka-Volterra model of interspecific competition:

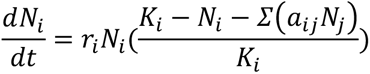

where *N_i_* is the mass of species *i*, *r_i_* is the maximal growth rate of species *i*, *K_i_* is the carrying capacity of species *i*, *a_i,j_* is the interaction coefficient between species *i* and *j*. To consider competition for a single resource within a fixed space, we define *a_ij_* as *r_i_*/*r_j_* and we scale the carrying capacity to 1, giving:

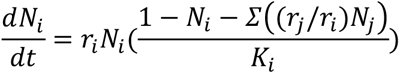

Additionally, we add a dilution term, δ_*i*_, to account for dispersal or removal from the system arriving at:

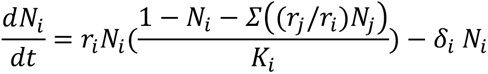

The coupling of the equations implements direct competition for resources, with the abundance of one species being limited by the abundance of the other. The coefficients for such competitive interactions are given by the relative growth between the species *r*_"_/*r*_*i*_, which favor species that grow faster. Dispersal is triggered when the *P. aeruginosa* population reaches an upper threshold *N*_*up*_ and is implemented by a dispersal rate δ_*i*_. Conversely, dispersal ceases when the *P. aeruginosa* population reaches a lower threshold *N*_%&’(_, restoring δ_*i*_ = 0. We simulate this system using MATLAB ODE solver *ode113* to integrate the system of equations over time. We simulate the system piecewise, changing the parameter δ_*i*_ whenever the *P. aeruginosa* population crosses the upper or lower thresholds.

We model movement of the different species across a 2D surface by adding a diffusion term, arriving at a reaction-diffusion model consisting of a system of partial differential equations:

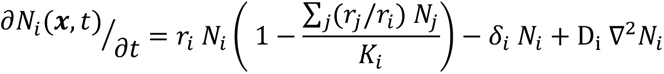

Dispersal is triggered locally when the local *P. aeruginosa* density reaches the upper threshold *N*_*up*_. We then simulate the model for an initial condition where the genotypes are randomly scattered across the surface. We use the Crank–Nicolson method, choosing time steps Δ*t* and distance increments *h* such that Δ*t* < 0.5 Δ*h*^2^/ *D* to guarantee the stability of the solution.

In our lattice-based model of interspecific competition, space is represented as a 2-dimensional discrete lattice sized 250×250 with periodic boundary conditions (the surface is a taurus). Each site can be vacant (state = 0) or occupied by a species of *i* (state=1,2,3). Individuals of type *i* are removed with probability *S_i_* and give birth with probability *β_i_*, and these probabilities are functions of the local environment. A type *i* born will try to occupy one of eight sites surrounding the seeding individual. If the chosen site is vacant, it changes to state *i*, otherwise nothing happens. When the simulation is run, at each time-step, sites are chosen at random to either replicate or become vacant. Movement to unoccupied grid sites is uniform for all three types.

The birth probability is set by the maximal growth rate for each species times the number of empty sites adjacent to each individual. The removal probability is set by the carrying capacity of each species times the number of occupied sites around each individual, rescaled to be the same order of magnitude as the maximal growth rate. This yields a single species stochastic-spatial model that is analogous to the ODE model,

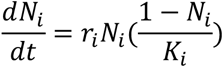

Since space is limited in the simulation to 250×250 grid nodes, competition occurs through the preferential occupation of free sites, without the need for the introduction of the *a*_*ij*_ parameter. For example, when a type dies it leaves a site open and whichever type can occupy that open site first will increase in abundance while the other type decreases in abundance. Increasing the maximal growth rate *r* gives an advantage in colonizing new sites while the population density is low. Increasing *K* gives an advantage in occupying vacant sites when the population density is high.

Mass dispersal events are implemented in a manner that matches our experimental data. At *N_up_* dispersal occurs as an increase in *P. aeruginosa* death probability, and *P. aeruginosa* sites rapidly become unoccupied. At *N_down_* dispersal stops and the death probability returns to normal. For all figures, we ran simulations with the same random seed. The MATLAB and Python scripts used to generate model figures are publicly available on GitHub and Zenodo. Parameters can be found in SI Table S2.

### Extended description of mathematical model results

We begin our theoretical exploration of the three dispersal scenarios, no dispersal, unbiased dispersal, and biased dispersal, with a mean field model, expand to a deterministic spatial model, and finally simulate a stochastic-spatial analogue of our mean-field model to capture key dynamic and spatial patterns found in our experimental system. While each model is parameterized using the same relative maximal growth rate and relative carrying capacity, and competition is implemented as a ratio of growth rates modified by carrying capacity (see Methods), they each have different assumptions and as a result give slightly different predictions of population dynamics and spatial patterning.

To follow the competing species in mixed communities over time, we model their population dynamics with coupled differential equations of logistic growth, reflecting competition for resources between the different genotypes, with additional terms for the dispersal of each species, which is triggered by high density of *P. aeruginosa* (see Methods). This mean-field model shows cycles of *P. aeruginosa* dispersal and recolonization, which allows coexistence of the three species when dispersal is biased (SI Figure S10A-C). Albeit, with a cyclic behavior that would likely result in the loss of one or more species in an environment with fixed population sizes and stochastic effects. In the biased dispersal condition, *E. coli* and *E. faecalis* take advantage of resources freed by *P. aeruginosa* dispersal to grow in numbers. We next determined the conditions for coexistence of multiple species by simulating the system under different levels of dispersal ratios, defined as the dispersal rate of *E. coli* and *E. faecalis* relative to *P. aeruginosa* (*δ_j_*/*δ_Pa_*), and different relative fitness between *P. aeruginosa* and its competitors, obtained by varying either the *P. aeruginosa* growth rate or its carrying capacity (Figure 2H, SI Figure S10D). We find that in the absence of dispersal, communities progress towards either *P. aeruginosa* dominance or elimination, depending on *P. aeruginosa*’s relative fitness. Increasing the level of dispersal bias, increases the window of *P. aeruginosa* relative fitness that permits three-species coexistence.

We then added spatial constraint to our mean-field model by assuming that cells can move through a surface, occupying new territory wherever nutrients are still available. We model this movement as diffusion without any directional bias, allowing cells to move freely from higher to lower density areas, resulting in a reaction-diffusion system that tracks the density of each genotype across a 2D space. Provided that the diffusion of cells through the surface is slow enough to retain some spatial structure, we now can have regions dominated by each species, with interactions happening mostly at the interfaces of these regions. Although the maximum growth rate of *P. aeruginosa* is slower than *E. faecalis’*s, it can reach higher local densities, which eventually results in dominance over the whole surface. As localized regions reach the threshold *P. aeruginosa* density and dispersal ensues, resources in these areas are then freed for the cells that remain. When dispersal is biased, these areas are preferentially occupied by *E. coli* and *E. faecalis.* When dispersal is unbiased, these areas are preferentially occupied by only *E. faecalis* (SI Figure S11).

The dispersal process creates regions with lower *P. aeruginosa* density, which can absorb cells migrating from regions with higher density and prevent the *P. aeruginosa* threshold density from being reached everywhere, resulting in a heterogeneous landscape that simultaneously harbors both low- and high-density regions. Analyzing the local coexistence between different species, we find that *E. coli* and *E. faecalis* preferentially occupies the areas with low density of *P. aeruginosa* and are displaced when *P. aeruginosa* densities are high (SI Figure S11D).

To account for stochastic and spatial effects simultaneously, we implement a stochastic-spatial analogue of the mean-field model. Our stochastic-spatial model implements discrete individuals interacting on a square grid, with periodic boundary conditions, in which local death and birth probabilities are functions of neighboring grid square occupancy and species composition (see Methods). As in the mean-field model, dispersal is triggered by high global density of *P. aeruginosa*. As in the previous models, biased dispersal of *P. aeruginosa* allows for coexistence of the three species (Figure 2A-C). However, in this model the spatial arrangement is also necessary for coexistence, even in the biased dispersal case, because the oscillations in the community, which are driven by *P. aeruginosa* dispersal and regrowth, combined with finite population sizes and random chance will lead to the loss of *E. faecalis* from the system unless there is some degree of spatial segregation (SI Figure S12D). Taken together, these simulations suggest that even a moderate bias towards *P. aeruginosa* dispersal, combined with some degree of spatial constraint, is sufficient to explain the coexistence of the three species observed within biofilms. These models support that the spatial constraint can either come from interacting patches as in the reaction-diffusion model or from the discrete localization of individuals within a focal environment as in the stochastic-spatial model. On a cursory level our data appears to support the focal environment model, but we do not collect sufficient experimental data to fully determine this question of pattern and scale.

